# Requirements for efficient endosomal escape by designed mini-proteins

**DOI:** 10.1101/2024.04.05.588336

**Authors:** Jonathan Giudice, Daniel D. Brauer, Madeline Zoltek, Angel L. Vázquez Maldonado, Mark Kelly, Alanna Schepartz

**Affiliations:** Department of Chemistry, University of California, Berkeley, CA 94720; Department of Molecular and Cellular Biology, University of California, Berkeley, CA 94720; School of Pharmacy, University of California-San Francisco, San Francisco, CA 94158; California Institute for Quantitative Biosciences (QB3), University of California, Berkeley, CA 94720; Chan Zuckerberg Biohub, San Francisco, CA 94158; Arc Institute, Palo Alto, CA

## Abstract

ZF5.3 is a compact, rationally designed mini-protein that escapes efficiently from the endosomes of multiple cell types. Despite its small size (27 amino acids), ZF5.3 can be isolated intact from the cytosol of treated cells and guides multiple classes of proteins into the cytosol and/or nucleus. In the best cases, delivery efficiencies reach or exceed 50% to establish nuclear or cytosolic concentrations of 500 nM or higher. But other than the requirement for unfoldable cargo and an intact HOPS complex, there is little known about how ZF5.3 traverses the limiting endocytic membrane. Here we delineate the attributes of ZF5.3 that enable efficient endosomal escape. We confirm that ZF5.3 is stable at pH values between 5.5 and 7.5, with no evidence of unfolding even at temperatures as high as 95 °C. The high-resolution NMR structure of ZF5.3 at pH 5.5, also reported here, shows a canonical p zinc-finger fold with the penta-arg motif integrated seamlessly into the C-terminal ⍺-helix. At lower pH, ZF5.3 unfolds cooperatively as judged by both circular dichroism and high-resolution NMR. Unfolding occurs upon protonation of a single Zn(II)-binding His side chain whose p*K*_a_ corresponds almost exactly to that of the late endosomal lumen. pH-induced unfolding is essential for endosomal escape, as a ZF5.3 analog that remains folded at pH 4.5 fails to efficiently reach the cytosol, despite high overall uptake. Finally, using reconstituted liposomes, we identify a high-affinity interaction of ZF5.3 with a specific lipid–BMP–that is selectively enriched in the inner leaflet of late endosomal membranes. This interaction is 10-fold stronger at low pH than neutral pH, providing a molecular picture for why escape occurs preferentially and in a HOPS-dependent manner from late endosomal compartments. The requirements for programmed endosomal escape identified here should aid and inform the design of proteins, peptidomimetics, and other macromolecules that reach cytosolic or nuclear targets intact and at therapeutically relevant concentrations.

---

To be effective, a potential therapeutic macromolecule must circumnavigate a complex path to traffic from the extracellular space into the cytosol or nucleus of a mammalian cell, and it must do so efficiently and without degradation of sequestration. Although a small number of natural or designed cyclic peptides can diffuse passively through biological membranes, these molecules are the exception, not the rule^1^. For most macromolecules, whether free or encased in various nanocarriers, the path into internal cell compartments such as the cytosol involves endocytosis, the natural process by which cells survey the extracellular environment. During endocytosis, material associated with the cell surface is packaged into membrane-delimited endosomes that mature through directional and biochemically distinct stages^2,3^. Release of material from endosomes into the cytosol without endosomal rupture requires passage of a macromolecule through a membrane that is not obviously any more permeant than the plasma membrane. Although it has been known for more than 35 years that certain transcription factors can activate transcription when added to cells in culture^4,5^, it has proven difficult to translate this finding into a robust strategy for therapeutic macromolecule delivery. Designing molecules that gain entry into the endocytic pathway is not the problem - the problem is endosomal escape.

ZF5.3 is a compact, rationally designed mini-protein that escapes from the endosomes of multiple cell types with high efficiency^6,7^. Despite its small size (27 amino acids), ZF5.3 can be isolated intact from the cytosol of treated cells^7^, and guides multiple classes of proteins, including enzymes^8–10^, transcription factors^11^, and monobodies^10,12^, into the cytosol and/or nucleus. In the best cases, delivery efficiencies reach or exceed 50% to establish nuclear or cytosolic concentrations of 500 nM or higher^7,10^. The egress of ZF5.3 from endosomes occurs through a biochemically distinct pathway that demands HOPS, a ubiquitous tethering complex that mediates endosomal maturation^13^. Egress of ZF5.3 and covalent ZF5.3-conjugates from endosomes proceeds without leakage of other intralumenal components^13^, and is especially efficient when the cargo protein is small, intrinsically disordered, or unfolds at a temperature of 35 °C or lower^10^. But other than the requirement for unfoldable cargo^10^ and an intact HOPS complex^13^, there is little known about how ZF5.3 traverses the limiting endocytic membrane. It is especially unclear why the endosomal membrane is favored over, say, the plasma membrane. Thus precisely what attributes of ZF5.3 enable efficient endosomal escape, and whether these attributes could be generalized, remain unclear.

Here we describe structural, biochemical, and chemical biology experiments that delineate the attributes of ZF5.3 that enable efficient endosomal escape. We confirm that ZF5.3 is stable at pH values higher than pH 5, with no evidence of unfolding at temperatures as high as 95 °C. The high-resolution NMR structure of ZF5.3, also reported here, shows a canonical ββ⍺ zinc-finger fold with the penta-arg motif integrated seamlessly into the C-terminal ⍺-helix. Although ZF5.3 is stably folded at neutral pH, it unfolds cooperatively at pH values between 4 and 5 as judged by both circular dichroism and high-resolution NMR. More detailed NMR studies reveal that unfolding occurs upon protonation of a single Zn(II)-binding His side chain whose p*K*_a_ (4.6) corresponds almost exactly to the pH of the late endosomal lumen. The unfolding of ZF5.3 is essential for endosomal escape, as an analog that remains folded at pH 4.5 also fails to efficiently reach the cytosol, despite efficient overall uptake. Finally, we discover a high-affinity interaction of ZF5.3 with a specific lipid–BMP–that is selectively enriched in the inner leaflet of late endosomal membranes. The interaction between ZF5.3 and BMP is substantially stronger at low pH than at neutral pH, providing a simple explanation for why escape occurs preferentially and in a HOPS-dependent manner from late endosomal compartments. The requirements for programmed endosomal escape identified here should aid the design of proteins, peptidomimetics, and other macromolecules that reach cytosolic or nuclear targets at therapeutically relevant concentrations.

## Results

### ZF5.3 unfolds cooperatively at the pH of the late endosomal lumen

Previous work has shown that the escape of ZF5.3 from the endosomal pathway occurs selectively from endolysosomes^13^, vesicular compartments whose lumenal pH is substantially lower than the bulk cytosol^14^. As canonical C_2_H_2_ zinc finger domains^15^ unfold at low pH, we first used circular dichroism (CD) spectroscopy to assess if this pH change would affect ZF5.3 secondary structure. ZF5.3 (**Fig. 1a**) used for CD analysis was generated recombinantly or using solid phase methods and purified to homogeneity (**Extended Data Fig. 1a,b**). The CD spectrum of ZF5.3 at pH 7.5 shows strong negative ellipticity at 208 nm, as expected for the ββ⍺ fold^17^ adopted by C_2_H_2_ zinc finger domains (**Fig. 1c**) and was unchanged as the concentration of ZF5.3 increased from 25 to 250 µM (**Extended Data Fig. 1c**). Temperature-dependent CD studies revealed that ZF5.3 is stable at pH 7.5, with no detectable change in ellipticity at 208 nm even at temperatures approaching 95 °C (**Fig. 1e**). To assess the effect of pH on ZF5.3 structure, we measured CD spectra at pH values between 3.5 and 7.5 (**Fig. 1b**). An overlay of these spectra reveals a shift of the major ellipticity minimum to lower wavelength as the pH decreases from 7.5 to 3.5. A plot of the change in ellipticity at 208 nm as a function of pH shows a cooperative transition with an apparent p*K*_a_ of 4.6 (**Fig. 1c**). Notably, we again detected no change in ellipticity at 208 nm when a pH 4.5 solution of ZF5.3 was heated to 95 °C (**Fig. 1e**). No pH-dependent changes in secondary structure were observed by CD when ZF5.3 was reconstituted in a Zn(II)-free buffer **(Fig 1d)**. We conclude that ZF5.3 undergoes a cooperative, pH- and Zn(II)-dependent change in structure with a pH midpoint of 4.6. This value corresponds almost precisely to the pH of the late endosomal lumen^14^.

**Fig 1.**
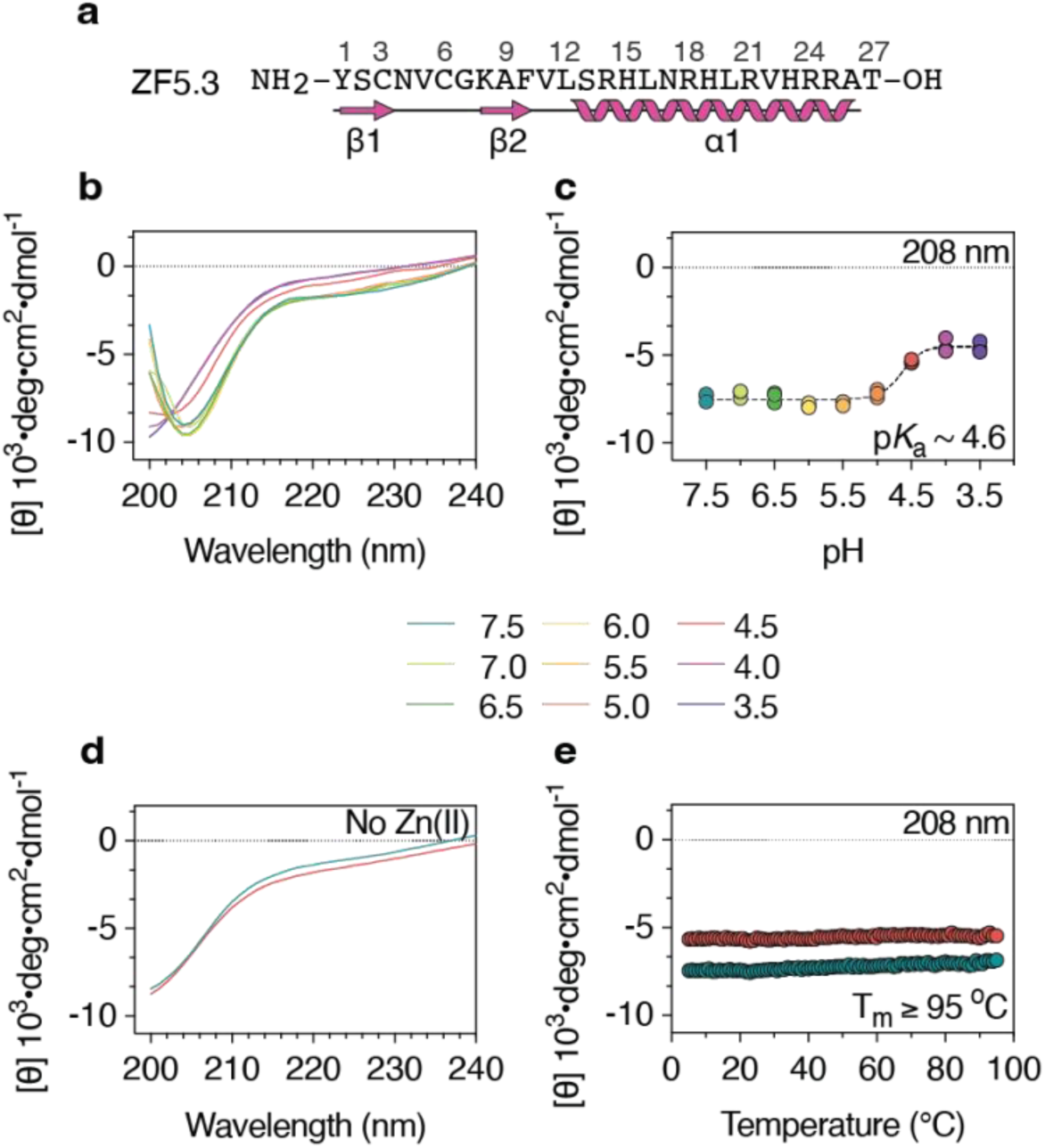
ZF5.3 unfolds cooperatively at the pH of the late endosomal lumen. **a**, The primary sequence of ZF5.3, annotated to identify regions of expected ⍺-helical and β-sheet secondary structure. Samples of ZF5.3 used for CD experiments carried free N- and C-termini and were generated either recombinantly (panels **b**, **c**, and **e**) or using solid-phase peptide synthesis (panel **d**) as described in **Methods**. CD experiments were performed at the indicated pH in a Resuspension Buffer containing 20 mM Tris-HCl, 150 mM KCl, 1 mM TCEP, and 100 µM ZnCl_2_. **b**, Smoothed wavelength spectra of 150 µM ZF5.3 monitored between 200 to 240 nm and at every 0.5 pH unit between pH 7.5 and 3.5 in Resuspension Buffer. **c**, Plot of the mean residue ellipticity at 208 nm as the pH of 150 µM ZF5.3 is titrated from pH 7.5 to pH 3.5. These data were fitted using a Boltzmann sigmoidal curve fit (R^2^ = 0.98) using GraphPad Prism 9 software. **d,** Wavelength spectra acquired for ZF5.3 in the absence of Zn(II) at pH 7.5 and pH 4.5. **e**, Plot of the mean residue ellipticity at 208 nm as 150 µM ZF5.3 at pH 7.5 (teal) or pH 4.5 (red) is heated from 2 °C to 95 °C every 1 °C.

### High-resolution NMR data provide evidence for pH-induced unfolding of ZF5.3

High-resolution NMR was used to learn more about the pH-dependent change in ZF5.3 structure detected by CD. Samples of ZF5.3 that were uniformly labeled with ^15^N or with both ^13^C and ^15^N were expressed in *E. coli*, purified to homogeneity, and reconstituted with Zn(II) (**Extended Data Fig. 2**). Initial experiments made use of ^15^N-labeled ZF5.3 and two dimensional [^15^N-^1^H] heteronuclear single quantum coherence spectroscopy (HSQC) to measure the chemical shifts of all nitrogens and their directly bonded protons as a function of pH^16^. At pH 5.5, the HSQC spectrum of ZF5.3 shows 24 discrete crosspeaks dispersed over a wide chemical shift range. These crosspeaks span from 6.5 to 9.3 ppm on the ^1^H axis and from 112 to 125 ppm on the ^15^N axis (**Fig. 2a**). Slightly fewer peaks are observed at pH 6.5 and 7.5, but the cross peaks observed are otherwise identical (**Extended Data Fig. 3**). When the pH is lowered to pH 5.0, the crosspeaks present at pH 5.5 remain but are now accompanied by a second set of 24 crosspeaks that are dispersed over a narrower range, between 7.0 and 8.6 on the ^1^H axis and between 116 and 125 ppm for all but three ^15^N resonances (**Fig. 2b**). When the pH is lowered once again to pH 4.5 and 3.5, the crosspeaks present at pH 5.5 disappear, and only the more narrowly distributed set of crosspeaks remain (**Fig. 2c, Extended Data Fig. 3**). The changes detected in the HSQC spectra of ZF5.3 as the pH is lowered systematically from 5.5 to 3.5 mirror those detected by CD, with a pH midpoint of approximately pH 4.5. This correlation is consistent with a cooperative, pH-dependent conformational change in ZF5.3 that alters the chemical environment of 24 of 27 amide NH protons.

**Fig. 2.**
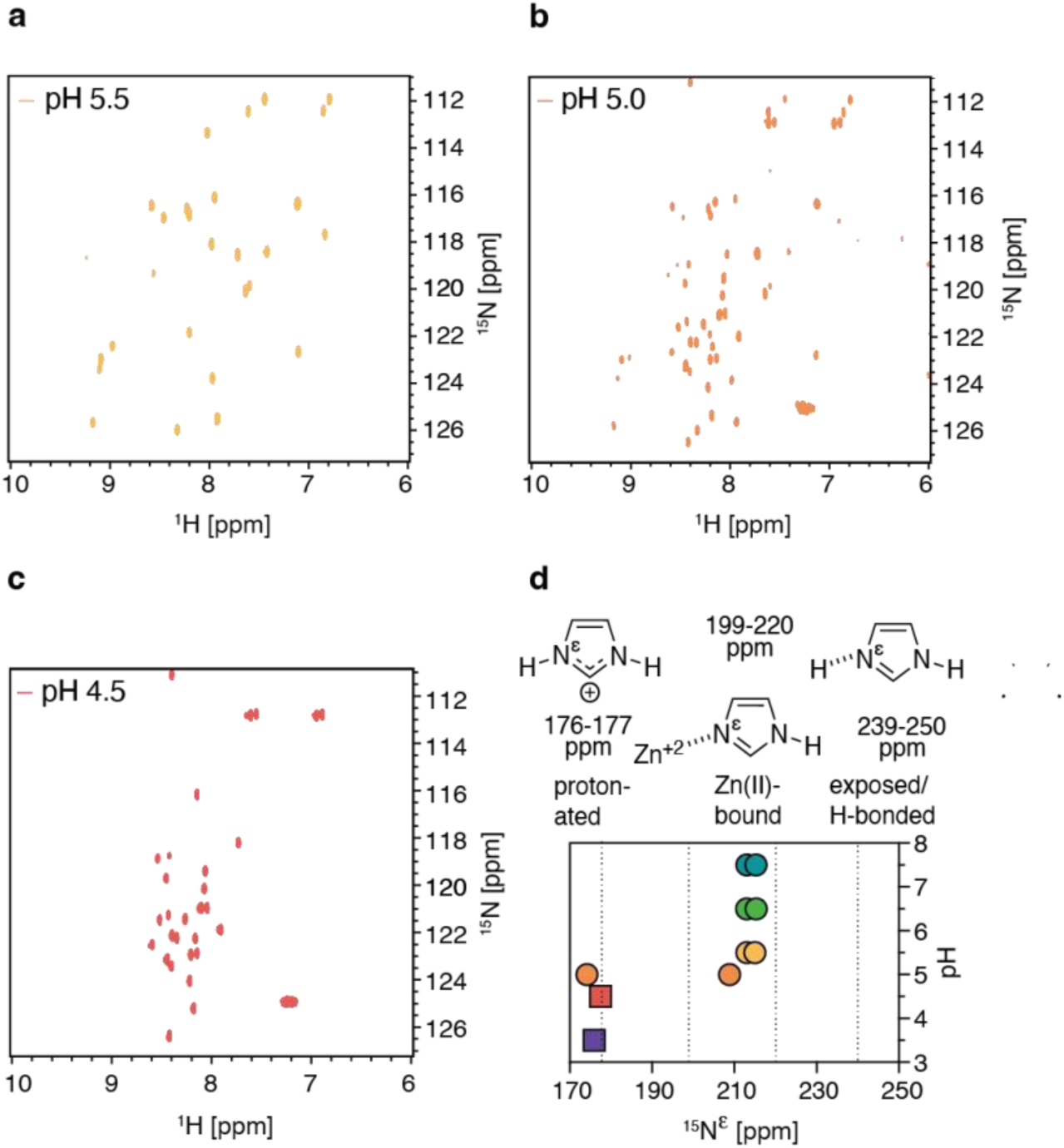
^15^N-^1^H heteronuclear quantum coherence spectroscopy (HSQC) data provide evidence for pH-induced unfolding of ZF5.3. **a-c**, ^15^N-^1^H HSQC of 800 µM ^15^N-labeled ZF5.3 acquired in a 20 mM citrate-phosphate buffer containing 10% D_2_O, 100 mM NaCl, 1 mM TCEP, and 1.6 mM ZnCl_2_ at pH 5.5, 5.0, and 4.5. d, Plot of the chemical shift of the N^ε^ in the two Zn(II)-coordinating His side chains of ZF5.3 (His19 and His23) at pH values between 3.5 and 7.5. At pH 5.0 and above, coordination of Zn(II) is observed for at least one His and represented on the plot as two circles, one for each Zn-bound His. At pH 4.5 and below all histidines are protonated with the same N^ε^ chemical shift and are represented on the plot as a single square.

### Unfolding of ZF5.3 at low pH results from protonation of a Zn(II)-bound His residue

The CD and NMR experiments presented above imply that a cooperative and Zn(II)-dependent change in ZF5.3 structure occurs between pH 5.0 and 4.5. The fact that this change is observed only for the Zn(II)-bound form of ZF5.3 and not for the Zn(II) free form (**Fig. 1d**) focused our attention on the effect of pH on the four residues within ZF5.3 that coordinate Zn(II) directly: Cys3, Cys6, His19, and His23. Zn(II)-bound His residues protonate in precisely the same pH-range as the pH-induced conformational changes within ZF5.3. Their chemical shifts vary with pH as well as with the state (bound *vs.* unbound) and location (N^ε^ *vs.* N^δ^) of Zn(II) coordination^17^. We hypothesized that protonation of a Zn(II)-bound His could trigger the pH-dependent conformational change detected by CD and NMR. To test this hypothesis, we made use of ^15^N-labeled ZF5.3 and long-range 2D [^15^N-^1^H]-HSQC experiments with an additional delay of 22 ms (1/(2JNH)) to refocus single-bond correlations and monitor the ^15^N^ε^ and ^15^N^δ^ chemical shifts of all three His residues within ZF5.3^18^.

Previous studies have shown that the ^15^N^ε^ chemical shift of a solvent exposed His side chain falls near 249.5 for a His β tautomer or 167.5 ppm for a His ɑ tautomer, near 239.5 for a His β tautomer or 177.5 ppm for a His ɑ tautomer when the N^ε^ participates in a hydrogen bond, between 199-220 ppm when the N^ε^ is Zn(II)-bound within a His β tautomer, and between 176 and 177 ppm when the N^ε^ is protonated regardless of tautomeric identity (**Fig. 2d**)^17,18^. The ^15^N^ε^ chemical shift falls within 170 - 177 ppm when the N^ε^ is Zn(II)-bound within an ɑ tautomer. We used triple-resonance CBCANH and CBCAcoNH experiments at pH 5.5, accompanied by 2D hbCBcgcdHD and 2D ^13^C Constant-Time HSQC experiments, to assign the two Zn(II)-coordinating His side chains – His19 and His23 – as β-tautomers and the non-Zn(II)-coordinating residue His15 as the ɑ-tautomer (**Extended Data Fig. 4**)^19,20^. Next, we monitored changes in the ^15^N^ε^ chemical shifts of His19 and His23 as the solution pH decreased from 7.5 to 3.5 (**Fig. 2d, Extended Data Fig. 5, Table 1**). No changes in the ^15^N^ε^ chemical shift of either residue were observed between pH 7.5 and 5.5; in this range both His residues display a ^15^N^ε^ chemical shift consistent with the Zn(II)-bound form (**Fig. 2d, Extended Data Fig. 5, Table 1**). At pH 5.0, however, the ^15^N^ε^ resonances of one His side chain falls in the range expected for a protonated side chain and at pH 4.5 and below, both ^15^N^ε^ resonances fall in this range. Thus, the changes detected in the ^15^N^ε^ chemical shift as the pH is lowered systematically from 5.5 to 3.5 mirror those detected by CD and in the HSQC spectra of ZF5.3. Taken together, these experiments support the hypothesis that the cooperative change in ZF5.3 structure that occurs at low pH results from protonation of one or both Zn(II)-bound His side chains. None of the three His side chains within ZF5.3 shows evidence of coordinating Zn(II) at pH values below 4.5 (**Fig. 2d**).

### At neutral pH, ZF5.3 assembles into a canonical CCHH zinc-finger fold

To better characterize the cooperative change in ZF5.3 structure induced at low pH, we first determined its structure at a pH above the low-pH transition. A series of 2D and 3D double and triple-resonance experiments performed at pH 5.5 were used to sequentially assign 89% of the ^1^H, ^15^N, and ^13^C backbone and side-chain resonances of ^13^C/^15^N-enriched ZF5.3. Sequential backbone assignments were performed manually using CCPNMR and validated using ARTINA^21–23^. Distance restraints were measured using a simultaneous 3D 13C/15N-edited NOESY-HSQC with a 120 ms mixing time. Dihedral angle restraints were predicted from NMR secondary chemical shifts using the neural network base program TALOS-N^24^. Structure determination utilized an AI assisted automatic NMR structure determination pipeline ARTINA^22^ implemented on the NMRtist web server^23^, which utilizes a trained deep neural network to identify signals in NMR spectra, resonance assignment using FLYA^25^ and automatic nOe assignments and automatic structure determination using CYANA^26^. The final NMR ensemble was calculated based on 392 nuclear Overhauser effect restraints (nOe; 90 long-range nOes (|i-j| > 5), 81 medium-range nOes (1 < |*i-j*| < 4), 100 sequential nOes (|i-j| = 1), and 121 intra-residue nOes (|i-j| < 1) (**Extended Data Table 2**) and the initial set of twenty lowest-energy ZF5.3 conformers superposes with an average pairwise backbone root mean square deviation (RMSD) of (for residues 1 to 27) of 0.8 Å and 1.34 Å for backbone and heavy atoms, respectively (**Fig. 3a**)^22^.

**Fig 3.**
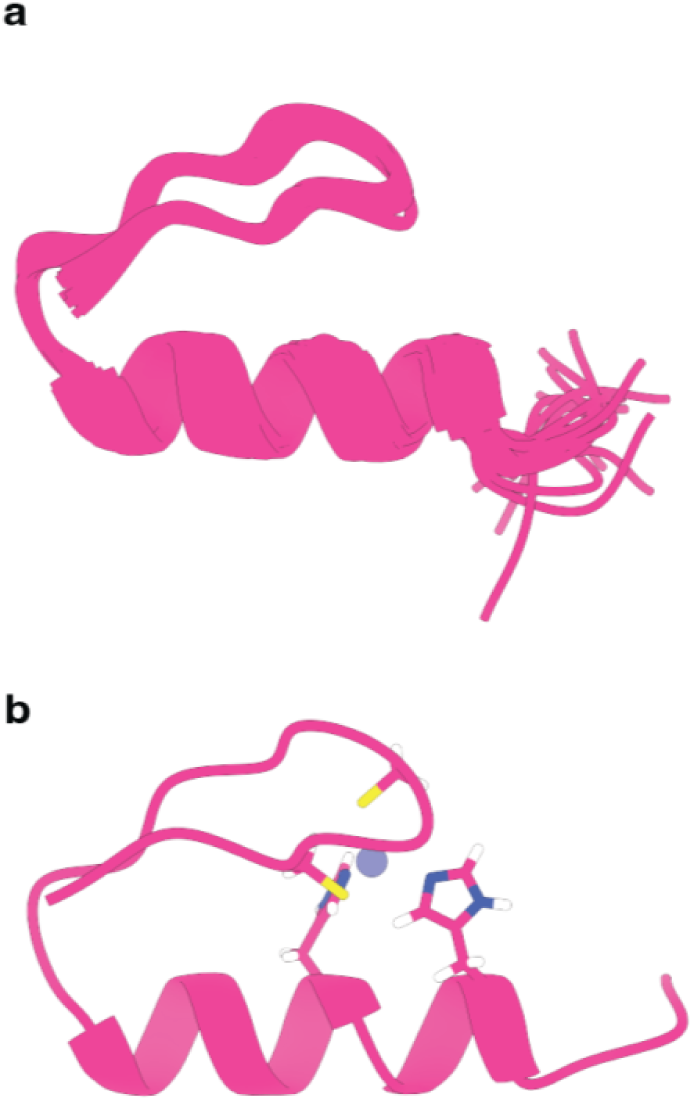
High-resolution NMR structure of ZF5.3 at pH 5.5. **a,** The initial set of twenty lowest-energy ZF5.3 conformers at pH 5.5 identified by ARTINA without specifying a discrete Zn(II) complex. **b,** Water-refined and minimized structure of Zn(II)-coordinated ZF5.3 (PDB: 9AZI).

We then made use of CYANA^27^ to introduce a single Zn(II) ion into the structural ensemble in a tetrahedral array coordinated by Cys3, Cys6, His19, and His23^26^. The structure was refined using restrained molecular dynamics and then minimized in explicit water using XPLOR-NIH v. 3.8 to provide a final ensemble of 20 Zn(II)-bound ZF5.3 conformers. These 20 conformers displayed a backbone RMSD to the mean of 1.2 Å (PDB accession code: 9AZI) (**Fig 3b**)^28,29^. The Zn(II)-bound ZF5.3 structure defined by this data comprises two short β-strands spanning residues Tyr1 - Cys3 and Lys8 - Phe10, a well-defined ⍺-helix spanning residues Ser13 - Arg24, and a flexible C-terminus containing amino acids Arg25 - Thr27. The ZF5.3 structural ensemble determined here aligns with that of human zinc finger protein 473 (PDB: 2EOZ), the canonical CCHH ZF from which ZF5.3 was designed, with a RMSD value of 0.899 ± 0.131 Å for superimposable atoms and an overall average RMSD value of 2.646 ± 0.875 Å upon Needleman-Wunsch structure alignment in ChimeraX (**Extended Data Fig. 6**)^30^. Taken together, these data indicate that at high pH ZF5.3 assembles into a canonical CCHH fold, despite the presence of a C-terminal penta-arginine motif^6^.

### Residue level characterization defines low pH conformer(s) of ZF5.3

As described previously, ^15^N-^1^H HSQC analysis of ZF5.3 at pH 4.5 indicates a substantial pH-dependent reduction in chemical shift dispersion across both the ^15^N and ^1^H axes compared to data obtained at pH 5.5 (**Fig. 2c**). This change is consistent with the population of less-or unstructured ZF5.3 conformers at low pH. To further define the low-pH conformation(s) of ZF5.3, we performed analogous 2D and 3D NMR experiments at pH 4.5. Over 90% of the amino acid resonances of ZF5.3 could be confidently assigned at pH 4.5. With both pH 5.5 and pH 4.5 ^15^N-^1^H HSQC spectra assigned, we calculated the pH-dependent Δδ (^15^N-^1^H) combined chemical shift perturbation^31^ (CSP) of each residue within ZF5.3 using the weighted absolute difference between the ^15^N and ^1^H chemical shifts of each residue at pH 4.5 and 5.5 (**Fig. 4a**). The greatest CSP values were associated with Val5 in the N-terminal β-strand (CSP = 1.62 ppm), the proximal ⍺-helix residue His23 which also coordinates Zn(II) (CSP = 1.79 ppm); and His15 and Arg21, which also residues in the C-terminal ⍺-helix (CSP = 1.61 and 1.57 ppm respectively). Moderate Δδ ^15^N-^1^H CSP values were observed for Phe10 and Leu16 (CSP = 0.69 and 1.14 ppm respectively), two hydrophobic residues implicated in maintenance of the conserved ββ⍺ fold, and for Cys3, Cy6, and His19 (CSP = 1.03, 0.74, and 0.95 respectively) the three remaining components of the Zn(II) chelate (**Fig. 4b**). The smallest CSP values were associated with Ala26 and Thr27, both of which are located near the ZF5.3 C-terminus (CSP = 0.09 and 0.03 ppm respectively). These Δδ (^15^N-^1^H) CSP values suggest that pH-induced unfolding involves the entire sequence of ZF5.3. In particular, the magnitude of the changes at residues Cys3, Cy6, His19, and His23 provides further evidence that the unfolded forms of ZF5.3 that exist at pH 4.5 lack a well-folded Zn(II) chelate.

**Fig 4.**
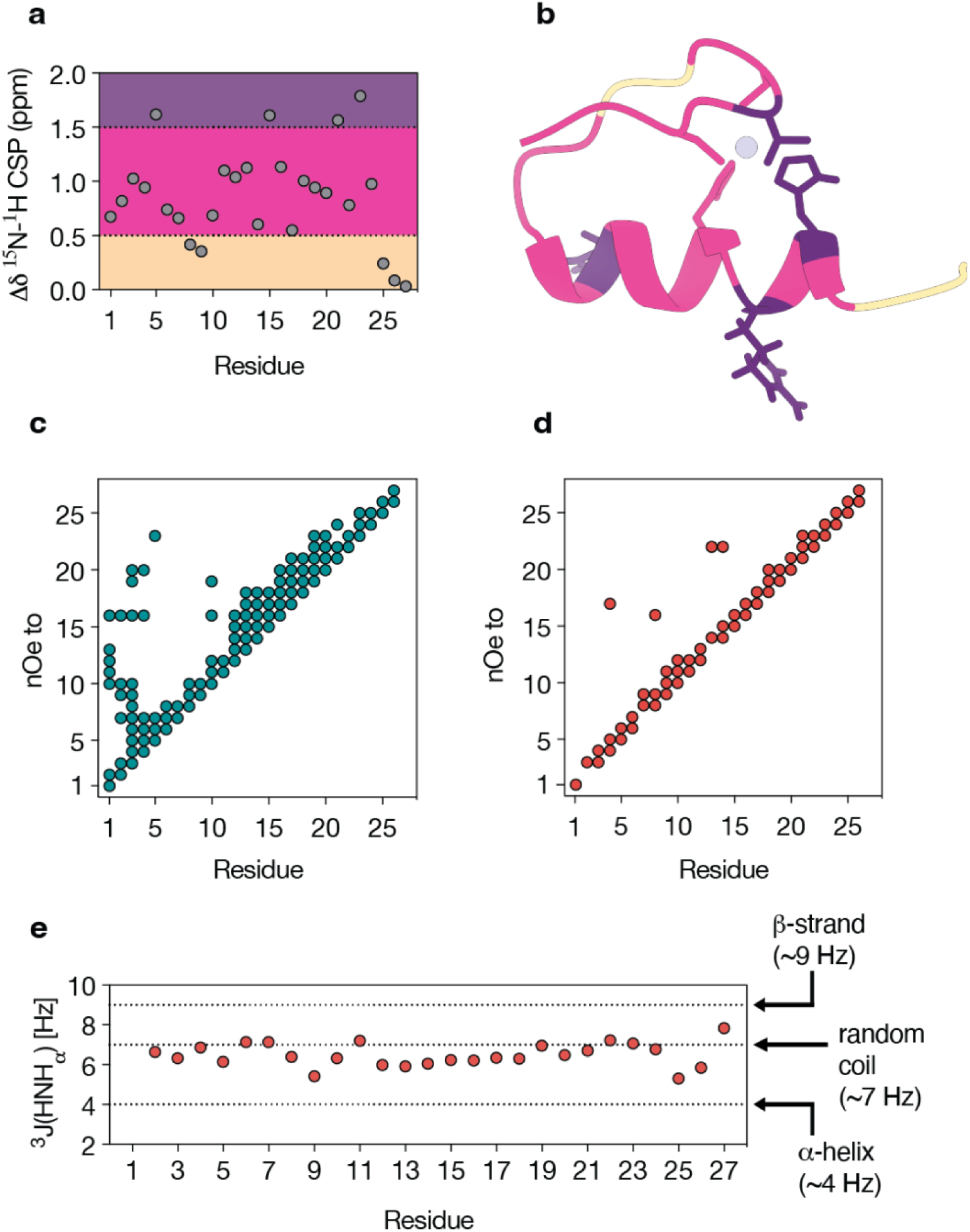
Residue level characterization defines low pH ZF5.3 conformer(s). **a,** combined ^15^N,^1^H CSP (*Δδ*) values quantify the weighted combined absolute difference in ^15^N and ^1^H chemical shift (*δ*) of each residue in ZF5.3 between pH 4.5 and 5.5 using the equation*Δδ* = 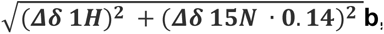, Image of the folded structure of ZF5.3 with each residue color-coded on the basis of the ^15^N,^1^H CSP (*Δδ*) value shown in panel a. **c,** nOe contact plot generated for ZF5.3 from restraints observed at pH 5.5 where each nOe is shown as a single point, regardless of whether multiple interactions were observed. **d,** nOe contact plot generated for ZF5.3 from restraints at pH 4.5. e, Plot of the *χ*1 side-chain torsional angle measured for each residue in ZF5.3 at pH 4.5 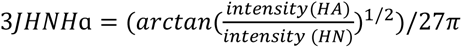 with reference to known values for amino acids located within specific secondary structural elements.

To examine whether any stable tertiary fold existed within ZF5.3 at pH 4.5, we measured nOe restraints using the same 3D ^13^C and ^15^N-edited NOESY-HSQC experiments used to generate the pH 5.5 ensemble, but in this case using dually labeled ^13^C and ^15^N ZF5.3 at pH 4.5. A total of 194 total nOes were identified in the pH 4.5 spectra: 13 long-range nOes (|*i-j*| > 5), 6 medium-range nOes (1 < |*i-j*| < 4), and 175 short-range nOes (|i-j| ≤ 1). The total number of nOes detected at pH 4.5 is roughly one-half of the 396 nOes (76 long-range, 82 medium-range, and 238 short-range) identified at pH 5.5. A side-by-side plot of the nOes detected within ZF5.3 at high and low pH highlights the significant loss of long-range restraints (**Fig. 4c**). The 13 long-range nOes detected at low pH were recorded between just 4 residues. Long-range nOes are detected between a residue within the N-terminal region of ZF5.3 that exists as a ꞵ-strand at high pH (Asn4 or Lys8) and a residue in the ⍺-helical region in the folded structure (Asn17 and Leu16, respectively) (**Fig. 4d**). Additional long-range nOes are detected between Val22 and both Ser13 and Arg14 (**Fig. 4d**). Not one of the long-range nOes detected in the low pH spectra of ZF5.3 were observed in NOESY spectra measured at higher pH, suggesting that the conformation(s) that are populated at low pH differ substantially from the structure determined at high pH.

To determine if residues in ZF5.3 experiencing long-range nOes at low pH could potentially reside within regions of local structure, we measured vicinal spin-spin three-bond HN-Hɑ J-coupling constants (^3^J_HNHɑ_) on a sample of ^15^N-labeled ZF5.3 at pH 4.5. The intervening *χ*1 side-chain dihedral angle unique to the secondary structure present locally within the residue can then be determined empirically. Specifically, ^3^J_HNHɑ_ values of 4 Hz are expected for residues in an ⍺-helical secondary structure, whereas ^3^J_HNHɑ_ values of 9 and 7 Hz are expected for residues within a β-strand or random coil, respectively^32^. Values for ^3^J_HNHɑ_ coupling constants were determined from ^15^N, ^1^H, and Hɑ chemical shifts measured in 3D HNHA spectra and quantified. A plot of the ^3^J_HNHɑ_ coupling constant calculated for each residue of ZF5.3 (residues 2 through 27) at pH 4.5 indicates that all residues experience *χ*1 side-chain torsional angles consistent with those found within a random coil (**Fig. 4e**). We conclude that the conformational landscape of ZF5.3 at pH 4.5–in aqueous solution–contains predominantly unfolded polypeptide chains that lack a discrete Zn(II) complex.

### Endosomal escape of ZF5.3 requires low pH-induced unfolding

Previous data indicate that ZF5.3 escapes efficiently from endolysosomes in a HOPS-dependent process^13^. CD and NMR data presented here indicate that ZF5.3 unfolds at the low pH associated with the endolysosomal lumen and lacks a Zn(II) chelate. But is unfolding of ZF5.3 at low pH necessary to move ZF5.3 across the endosomal membrane? And if so, what is the driving force for this translocation? And why is HOPS required? To address the first of these questions, we sought an analog of ZF5.3 that would retain the ability to gain entry into the endocytic pathway but simultaneously resist unfolding within the low pH endolysosomal lumen. As we postulate that the protonation of His23, a Zn(II)-bound residue, drives pH-induced ZF5.3 unfolding, we sought a “zinc-free” zinc finger that would resist low pH-induced unfolding and retains the canonical ββ⍺ fold at all pH values between 4 and 8.

Many years ago, Imperiali and coworkers designed a series of “zinc-free” zinc finger domains whose ββ⍺ fold is stabilized not by Zn(II) coordination but instead by hydrophobic interactions^33^. In particular, the 23-residue BBA5 folds into a structure that closely mirrors that of ZF5.3. The pH 5.5 structure of ZF5.3 determined here (**Fig. 3**) aligns with that of BBA5 (PDB: 1T8J) with an average RMSD value of 0.899 ± 0.092 Å for superimposable atoms and an overall average RMSD value of 3.949 ± 0.208 Å upon Needleman-Wunsch structure alignment in ChimeraX – the greatest differences are in the ꞵ-sheet region, as expected (**Fig. 5b**)^30^. BBA5.3 was generated from BBA5 simply by integrating the requisite penta-Arg motif from ZF5.3 into the BBA5 ⍺-helix with the proper alignment and spacing (**Fig. 5a**). The four residues of BBA5 that are essential for the Zn-free ββ⍺ fold (Tyr6, Phe8, Leu14, and Leu17) were not altered.

**Fig 5.**
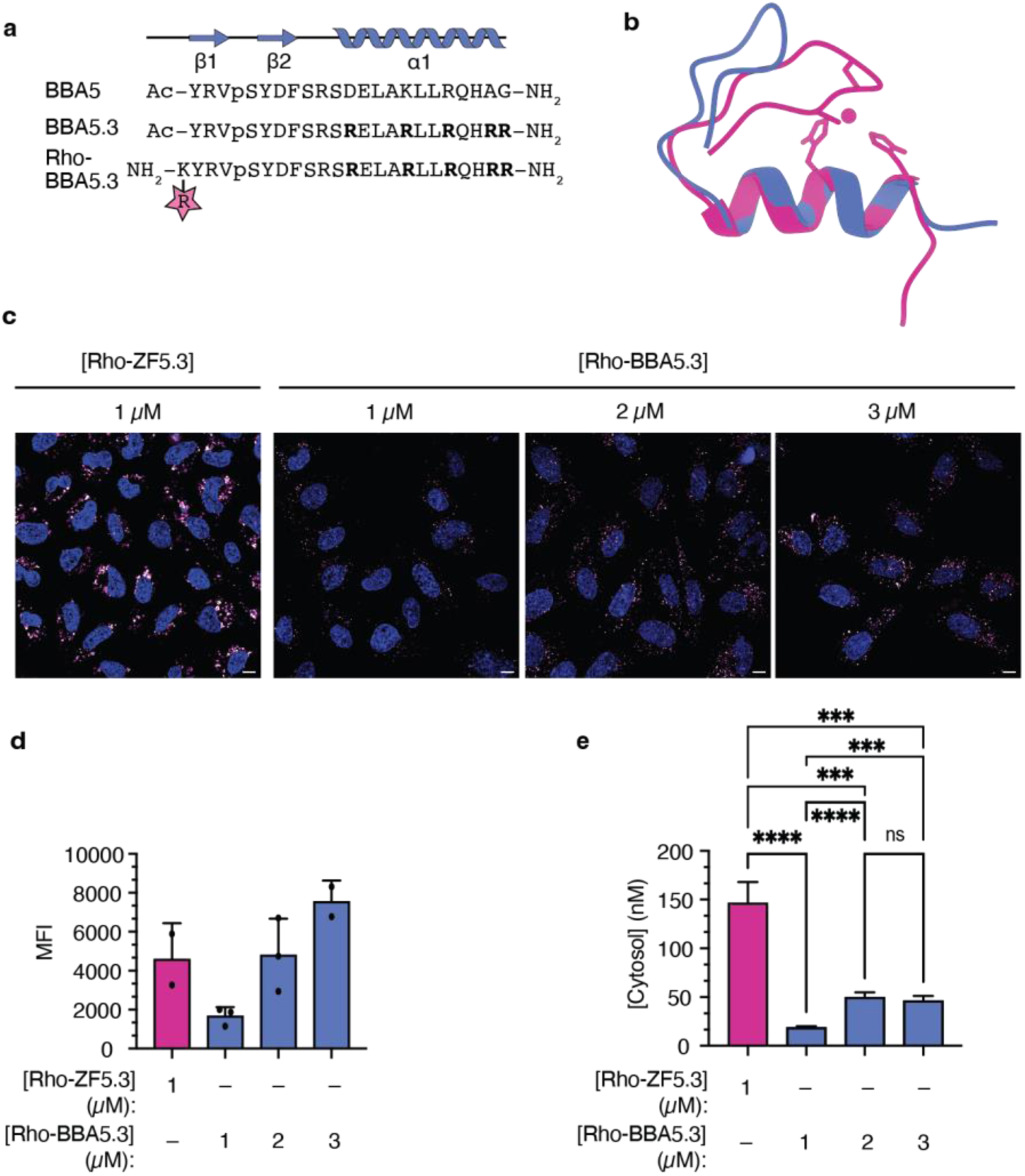
The pH-insensitive ZF5.3 variant BBA5.3 fails to efficiently reach the cytosol. **a**, The primary sequences of BBA5, BBA5.3, and Rho-BBA5.3, annotated to identify regions of expected ⍺-helical and β-sheet secondary structure. The five Arg residues that comprise the penta-arginine motif are highlighted in bold. Rhodamine was attached covalently to BBA5.3 upon reaction of an *N*-terminal lysine side chain with Lissamine Rhodamine B as described in **Methods**. A lowercase p in the primary sequence denotes D-Proline. **b,** Overlay of the high pH (pH 5.5) structure of ZF5.3 (pink) and BBA5 (PDB 1T8J) (blue) created using ChimeraX^30^. **c,** Confocal microscopy of Saos-2 cells treated for 1 h with the indicated concentration of Rho-ZF5.3 or Rho-BBA5.3 (magenta) and 300 nM Hoechst 33342 (blue) to identify nuclei. Scale bar indicates 10 µm. **d,** Flow cytometry of values are provided as Median Fluorescence Intensity (MFI) detected between 570 and 602 nm, n = 10000 cells in total per condition containing at least 2 biological replicates each (mean ± SEM). **e**, Results of FCS analysis showing the average cytosolic concentration (in nM) established upon incubation of Saos-2 cells with the indicated concentration of Rho-ZF5.3 or Rho-BBA5.3 for 1 h, n > 15 cells for each FCS condition with two biological replicates each (mean ± SEM). Statistical significance comparing the given concentrations was assessed using the Brown-Forsythe and Welch one-way analysis of variance (ANOVA) followed by an unpaired t-test with Welch’s correction. ****p ≤ 0.0001, ***p ≤ 0.001, **p ≤ 0.01, *p ≤ 0.05. BBA5.3 and Rho-BBA5.3 were synthesized using solid phase methods and purified to homogeneity (**Extended Data** Fig 8).

The CD spectrum of 125 µM BBA5.3 at pH 7.5 showed strong negative ellipticity at 208 and 222 nm and was unchanged as the pH of the solution was lowered progressively from pH 7.5 to pH 4.5 (**Fig. 5b**). Confocal microscopy images of Saos-2 cells treated with Rho-BBA5.3 show evidence of punctate fluorescence, consistent with uptake into the endocytic pathway (**Fig. 5c**). Flow cytometry of these cells after 1 h incubation reveals that the overall uptake of Rho-BBA5.3 is roughly one-half that of an equivalent concentration of Rho-ZF5.3 (**Fig. 5d**). FCS, however, which specifically measures concentration, reveals that delivery of Rho-BBA5.3 to the cytosol is almost one-tenth as efficient as the delivery of an equivalent concentration of Rho-ZF5.3: when Saos-2 cells are treated with 1 µM Rho-BBA5.3, the fraction that reaches the cytosol is 8-fold lower than the fraction of Rho-ZF5.3 that reaches the cytosol under identical conditions (**Fig. 5e**). Even when added to cells at a concentration of 3 µM, where the total uptake of Rho-BBA5.3 surpasses the uptake of 1 µM Rho-ZF5.3, the amount of Rho-BBA5.3 that reaches the cytosol is 3-fold lower than that of ZF5.3 (**Fig. 5e**). Thus, although Rho-BBA5.3 contains a penta-arg motif and reaches the endocytic pathway, it traffics poorly across the endosomal membrane into the cytosol. These data indicate that the penta-arg motif alone is not sufficient for highly efficient delivery of ZF5.3 to the cytosol. The impaired delivery of Rho-BBA5.3 to the cytosol supports the hypothesis that low pH-induced unfolding of ZF5.3 is required for endosomal escape.

### ZF5.3 interacts with vesicles enriched in bis(monoacylglycero)phosphate (BMP) lipids

In this work we have shown that ZF5.3 unfolds at the pH associated with the late endosomal lumen (**Fig. 1**), existing at this pH as an ensemble of unfolded polypeptides that lack a discrete Zn(II) chelate (**Fig. 4**), and that the ability to unfold at low pH is a prerequisite for efficient endosomal escape (**Fig. 5**). Previous microscopy data indicate that ZF5.3 localizes within both the lumen and membrane of endolysosomes and intraluminal vesicles^13^ but shed no light on what guides ZF5.3 into the limiting membranes of these structures or what factors might stimulate HOPS-mediated endosomal escape.

Recomposition of the limiting endosomal membrane is a hallmark of endosomal maturation^34^. While the plasma membrane is composed largely of phosphatidylcholine (PC), phosphatidylethanolamine (PE), and cholesterol, the membranes of endolysosomes and intraluminal vesicles are depleted in cholesterol and enriched in phosphatidylinositols in the outer leaflet and the endosome-specific lipid bis(monoacylglycero)phosphate (BMP) in the inner leaflet^35,36^. BMP is selectively enriched in late endosomal and lysosomal compartments and plays a crucial role in regulating their function and dynamics. BMP interacts with Apoptosis Linked Gene 2 Interacting Protein X (ALIX) to regulate cholesterol storage in endosomes^37^, promotes intraluminal vesicle budding^38^, and binds to Hsp70 to prevent lysosomal permeabilization in cell stress. BMP is also required for the infectivity of vesicular stomatitis virus and dengue virus due to its role in facilitating nucleocapsid membrane fusion and release^38,39^. Given this precedent, we asked whether the association of unfolded ZF5.3 with BMP in the limiting endolysosomal membrane is required for efficient endosomal escape of ZF5.3, and if there were differences in the behavior of ZF5.3 and BBA5.3 that shed further light on the rules and interactions that promote efficient endosomal escape.

To investigate membrane interactions *in vitro,* we prepared large unilamellar vesicles (LUVs) whose membranes were composed of lipid mixtures mimicking those of the plasma membrane (PM), BMP-containing LAMP^+^ late endolysosomes or lysosomes (BMP+LE/LY), or BMP-rich intraluminal vesicles (ILV). We also prepared LUVs whose membranes mimicked those of LE/LYs but were depleted of BMP (BMP-LE/LY)^40,41^. Liposome formation was verified *via* dynamic light scattering (**Extended Data** Fig. 8a), and a fluorescent co-sedimentation assay evaluated the extent to which ZF5.3 interacts with each type of LUV^42^ (**Fig. 6a**) Experiments were performed using both Rho-ZF5.3 and Rho-BBA5.3 and at both neutral pH (7.5) and acidic (4.5) pH to monitor interactions with folded (at pH 7.5) and unfolded (at pH 4.5) forms of ZF5.3. Rho-ZF5.3 or Rho-BBA5.3 were co-incubated separately with each of the four LUV populations at a 1:100 (mol/mol) protein:lipid ratio (5 µM mini-protein:500 µM lipid) for 30 min and the LUVs were pelleted *via* ultracentrifugation. The fraction of Rho-ZF5.3 or Rho-BBA5.3 associated with each LUV was determined by dividing the fluorescence intensity of the resuspended pellet (containing LUV-associated Rho-ZF5.3 or Rho-BBA5.3) by the total fluorescence (supernatant + resuspended pellet). The sedimentation of Rho-ZF5.3 or Rho-BBA5.3 in the absence of LUVs was subtracted as background.

**Fig 6.**
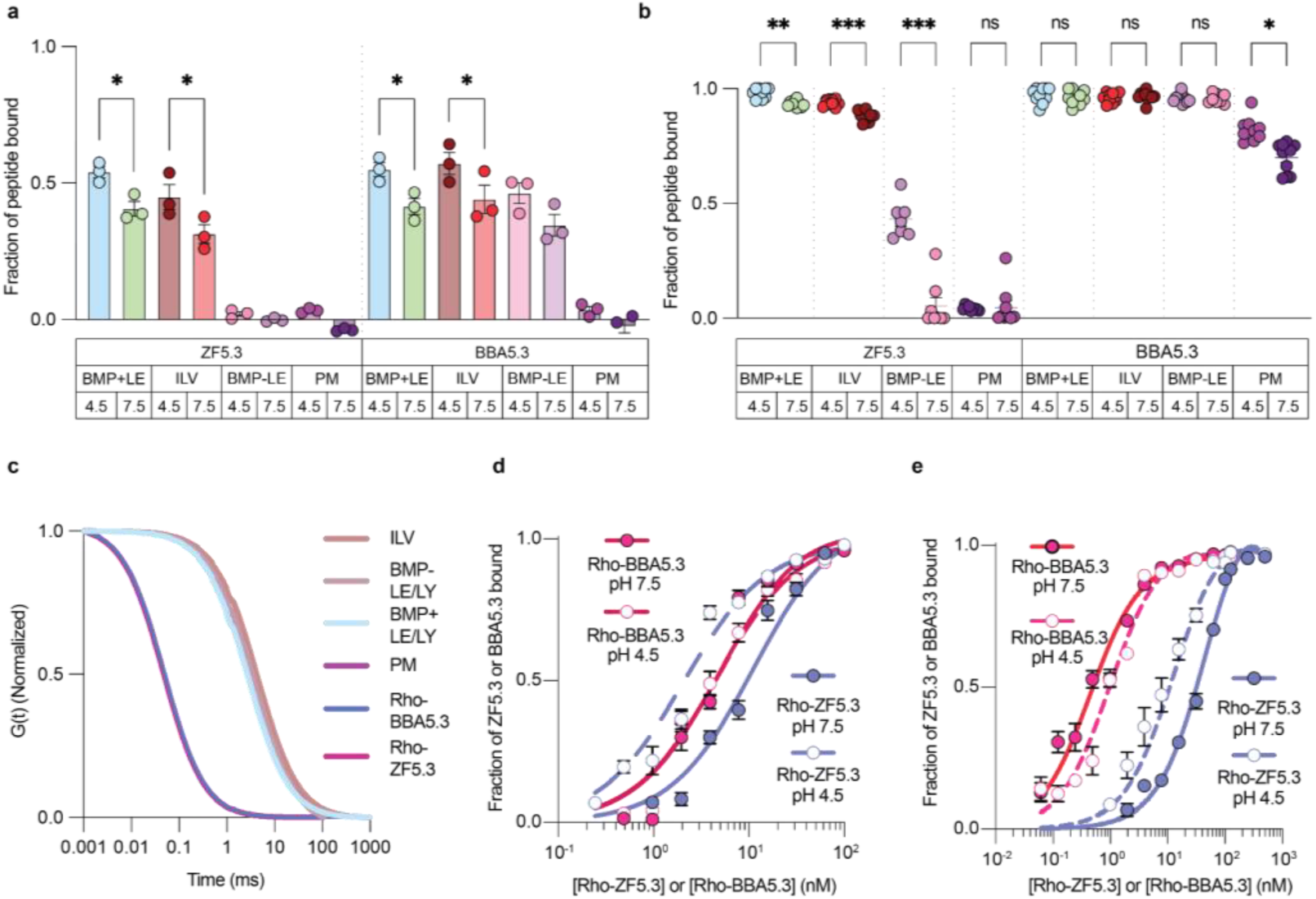
ZF5.3 displays pH-dependent binding to large unilamellar vesicles (LUVs) enriched in the late endosomal lipid bis(monoacylglycero)phosphate (BMP). **a**, Fractional binding of Rho-ZF5.3 and Rho-BBA5.3 to LUVs at pH 4.5 and 7.5 determined by fluorescent liposome sedimentation assay. Fractional binding (mean ± SEM) was determined as (pellet fluorescence intensity/total fluorescence intensity), n=3. Sedimentation of the mini-protein in the absence of LUVs was subtracted as background. Statistical significance comparing each condition at pH 4.5 vs pH 7.5 was assessed using the one-way analysis of variance (ANOVA) followed by Šídák’s multiple comparisons test, *p ≤ 0.05. **b,** Normalized autocorrelation curves of mini-proteins or LUVs of interest by *in vitro* single-component fluorescence correlation spectroscopy (FCS) (mean ± SEM), n=10. **c,** Fractional binding of Rho-ZF5.3 and Rho-BBA5.3 to LUVs at pH 4.5 and 7.5 determined by two-component FCS (mean ± SEM), n=10. Statistical significance comparing each condition was assessed using the Brown-Forsythe and Welch one-way analysis of variance (ANOVA) followed by Dunnett’s T3 multiple comparisons test. ****p ≤ 0.0001, ***p ≤ 0.001, **p ≤ 0.01, *p ≤ 0.05. **d,** Equilibrium binding curves of BMP+LE/LY LUVs and **e)** ILV LUVs to mini-proteins Rho-ZF5.3 and Rho-BBA5.3 at pH 4.5 and 7.5. Each point represents the mean ± SEM of the bound fraction determined by two-component FCS, n=10.

This fluorescent co-sedimentation assay revealed that the extent of LUV association depended on lipid composition and pH and in one specific case differed significantly between ZF5.3 and BBA5.3. Under these conditions (5 µM mini-protein:500 µM lipid), neither Rho-ZF5.3 nor Rho-BBA5.3 interacted substantially with LUVs whose phosphatidylcholine-rich composition mimics the plasma membrane, regardless of pH. In contrast, both Rho-ZF5.3 and Rho-BBA5.3 interacted substantially with BMP-rich LUVs that mimic the membranes of late endosomes and lysosomes (BMP+LE/LY LUVs), and in both cases the association was higher at pH 4.5 than at pH 7.5 (54 ± 2% and 41 ± 3%, respectively for Rho-ZF5.3; 55 ± 3% and 41 ± 3% for Rho-BBA5.3). The same trend held for LUVs that mimicked the composition of ILVs, whose membranes contain even higher BMP levels^35^. The biggest difference in the LUV-association of Rho-ZF5.3 and Rho-BBA5.3 related to the presence of BMP. The fraction of Rho-ZF5.3 associated with BMP-LE/LY LUVs was close to background regardless of pH, whereas the fraction of Rho-BBA5.3 associated with these LUVs was high at both pH 4.5 and 7.5 (46 ± 4% and 35 ± 4%, respectively) (**Fig. 6a**). These results imply that ZF5.3 and BBA5.3 interact with membranes whose lipids mimic late endolysosomes, and that ZF5.3 does so in a BMP-dependent fashion.

We used fluorescence correlation spectroscopy (FCS), a single-molecule method, to study LUV interactions at lower mini-protein concentration and a lower protein:lipid ratio to better resolve differences between the interactions of ZF5.3 and BBA5.3 (100 nM Rho-ZF5.3 or Rho-BBA5.3; 500 µM total lipid; 1:5000 (mol/mol) protein:lipid ratio). FCS can quantify the interaction of a fluorescently labeled protein or peptide with a lipid vesicle because of the large association-induced shift in the diffusion coefficient of the fluorescently tagged material^43^. First we established the *in vitro* diffusion coefficients of Rho-ZF5.3 and Rho-BBA5.3 in isolation, as well as those of each of the four LUV populations, which were labeled by the addition of 0.01% (mol/mol) 1,2-dioleoyl-sn-glycero-3-phosphoethanolamine-N-(lissamine rhodamine B sulfonyl) (Rho-PE) (**Fig. 6b**). The in vitro diffusion coefficients of Rho-ZF5.3 and Rho-BBA5.3 (261 ± 4 and 250 ± 7 µm^2^s^-1^, respectively) compared well to each other and to values previously reported for Rho-ZF5.3 (262 ± 3 µm^2^s^-1^). The in vitro diffusion coefficients of the PM, ILV, BMP+ LE/LY, and BMP-LE/LY LUVs were also similar to each other (3.7 ± 0.1, 2.7 ± 0.2, 3.0 ± 0.1, 3.8 ± 0.6, µm^2^s^-1^, respectively), consistent with reported values for 100 nm LUVs (1.4-5.8 µm^2^s^-1^), and notably ∼70-fold lower than the values for Rho-ZF5.3 or Rho-BBA5.3^13,44,45^.

To monitor the interaction of Rho-ZF5.3 and Rho-BBA5.3 with each LUV population, we mixed the components in 20 mM Tris pH 4.5 or 7.5, equilibrated at 37 °C for 30 min, and analyzed the resulting mixture by FCS. The resultant autocorrelation curves were fitted to a two-component function, where 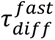is the diffusion time of free Rho-ZF5.3 or Rho-BBA5.3 and 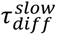 is the diffusion time of Rho-ZF5.3 or Rho-BBA5.3 bound to a LUV (**Extended Data Fig. 8b-e**). The fractional amplitude of each component of the fitting function provides the fraction of Rho-ZF5.3 or Rho-BBA5.3 bound to each LUV population (**Methods Equation 3**).

FCS analysis of LUV interactions with Rho-ZF5.3 and Rho-BBA5.3 largely recapitulated data obtained by the co-sedimentation assay but also identified low affinity interactions that were not detected previously. FCS recapitulated the substantial and pH-dependent association of Rho-ZF5.3 with BMP-rich LUVs that mimic endolysosomes (BMP+LE/LY) and intralumenal vesicles (ILV). It recapitulated the absence of pH-dependence for the association of Rho-BBA5.3 with BMP-rich LUVs that mimic endolysosomes (BMP+LE/LY) and intralumenal vesicles (ILV). It also recapitulated the dependence of Rho-ZF5.3, but not Rho-BBA5.3, on the presence of BMP. We do note that under these conditions, FCS also detected lower affinity and pH-dependent interactions of Rho-ZF5.3 with late endosome/lysosome-like LUVs that lacked BMP (BMP-LE/LY LUVs) and a lower affinity interaction of Rho-BBA5.3 with LUVs that mimicked the plasma membrane (**Fig. 6c**).

To quantify the extent to which pH influenced the equilibrium affinity of ZF5.3 for BMP+LE/LY LUVs and ILV LUVs, we acquired FCS autocorrelation curves using 60 pM −500 nM Rho-ZF5.3 in the presence of excess lipid (500 µM). Side-by-side experiments were performed with Rho-BBA5.3. Binding curves were fitted using non-linear regression to a total site binding model to determine an apparent equilibrium dissociation constant (*K*_d_) describing each association. The apparent *K*_d_ values describing the interaction of Rho-ZF5.3 with BMP+LE/LY LUVs at pH 4.5 and 7.5 are 2.1 nM and 11.3 nM, respectively and 12.6 nM and 52.8 nM, respectively for ILV LUVs. Thus in both cases, the interaction of Rho-ZF5.3 with BMP+LE/LY and ILV LUVs is more favorable at low pH than at high pH. This difference is not observed for Rho-BBA5.3. The equilibrium *K*_d_ values calculated for Rho-BBA5.3 at pH 4.5 and 7.5 are 973 pM and 446 pM, respectively for ILV LUVs and are 4.60 nM and 4.97 nM for BMP+LE/LY LUVs (**Fig. 6d-e**). In addition, BBA5.3 shows a lower *K*_d_ for ILV LUVs, which have a higher BMP content (77%) in comparison to BMP+LE/LY LUVs (20%) whereas ZF5.3 *K*_d_ is higher in ILV LUVs than in BMP+LE/LY LUVs. These values suggest that the membrane interactions of ZF5.3 may be inhibited by very high BMP content. Nevertheless, what is most notable is the 4-5-fold decrease in the affinity of Rho-ZF5.3 for BMP-containing LUVs upon increasing pH from 4.5 to 7.5. This pH gradient is precisely what ZF5.3 could experience during the process of HOPS-mediated membrane fusion.

## Discussion

Nature has devised several evolutionarily conserved strategies for moving proteins across otherwise impermeant barriers. A large number of proteins cross the limiting membranes of prokaryotic or eukaryotic cells in an energy-dependent process that requires a discrete translocon, a membrane-embedded protein or complex possessing an often hydrophilic channel that facilitates passage of an entire protein from one cellular compartment to another^46^. Some translocons convey folded proteins, while others convey proteins that are partially or completely unfolded. In addition, the structures of certain macrocyclic natural products, such as cyclosporine^47^, FK-506^48^, and griselimycin^49^, have evolved to cross impermeant barriers in a manner that demands neither ATP nor a discrete translocon^50^. Evidence suggests that in certain cases translocation is dependent on the unique ability to shape-shift between two conformations, one favored in an environment with high dielectric constant and another favored in the low dielectric constant that characterizes cellular membranes^47^.

The work reported here suggests that ZF5.3 escapes endosomes via a pathway that resembles in many respects that of cell-permeant macrocyclic natural products, but is selective for membranes that contain BMP and undergo vesicle fusion. In this work, we show that the efficiency of endosomal escape is dependent on cooperative unfolding of ZF5.3 at the pH associated with the late endosomal lumen. Using NMR, we show that pH-induced unfolding is a direct result of protonation of one or both of the His residues that coordinate Zn(II). BBA5.3, whose pI, overall fold, and size mimic that of ZF5.3, but lacks a zinc-chelate and the associated pH-induced unfolding transition, reaches endosomes efficiently but fails to traffic across these endosomal barriers into the cytosol. We further demonstrate the pH-dependent association of ZF5.3 with an anionic lipid that is selectively enriched in the inner leaflet of endosomal membranes. Perhaps most intriguing is the pH-dependence of the interaction of ZF5.3 with BMP-enriched liposomes, a pH-dependence that is not observed for BBA5.3, despite an identical pI.

Taken together, these features suggest a mechanism in which ZF5.3 preferentially associates with the limiting membrane of late endosomes that feature a lowered lumenal pH and a BMP-enriched membrane. Low pH-triggered unfolding of ZF5.3 may lower the energetic barrier for membrane crossing in comparison to a folded mini-protein such as BBA5.3. The data reported here suggest the ZF5.3 also undergoes translocation by virtue of a shape-shift, although in this case it is induced by a change in pH as opposed to a change in dielectric constant. It is possible that HOPS stimulates endosomal escape of ZF5.3 because HOPS stimulates endosomal acidification, as reported^51^; alternatively, HOPS may be involved directly in the escape process by facilitating membrane fusion between endosomal compartments in which ZF5.3 is unfolded and interacting with BMP at the membrane. And regardless of the role of HOPS, the involvement of an inadvertent yet essential translocon for ZF5.3 has not been rigorously excluded.

Although it has been known for more than three decades that certain transcription factors can reach the nucleus and function as gene regulatory agents when added to cells in culture^4,5^, the molecular mechanisms that support this activity have remained elusive. As a result, it has proven challenging to develop strategies for direct macromolecule delivery that engender more than a several-fold improvement in delivery efficiency. Short, cationic peptides that may possess a pH-dependent affinity with BMP-rich endosomal membranes are likely to be degraded or sequestered before they reach endolysosomes. Larger cationic proteins may enter the endocytic pathway and resist proteolysis, but their ability to escape from endosomes is impeded by the inability to gain access to the hydrophobic membrane without unfolding. Small protein domains that unfold in endosomes may not possess the sequence elements required for interactions with BMP and/or the as-yet-unidentified translocon. Thus, the complexity of the requirements for efficient endosomal escape by ZF5.3 provides a retrospective explanation for the challenges faced by previous direct macromolecule delivery strategies. Looking forward, the data presented here, alongside related work demonstrating that the optimal cargoes for ZF5.3 are those that are intrinsically disordered or easily unfold^10^, should aid in the design of proteins, peptidomimetics, and other macromolecules that reach cytosolic or nuclear targets at therapeutically relevant concentrations.

## Methods

### Synthesis of ZF5.3-OH (1)

ZF5.3 (**1**) was synthesized using Wang resin 100-200 mesh (Novabiochem) preloaded with Fmoc-Thr-OH at a 0.1 mmol scale using a GPT PurePep Chorus automated peptide synthesizer. The resin was first swelled for 20 min with DMF at RT. All Fmoc deprotection steps were performed twice using 5 mL of 20% 4-methyl piperidine in DMF at 50 °C for 2 min. Subsequent peptide coupling reactions were performed by delivering 5 equiv. of the requisite Fmoc-protected ⍺-amino acid (Novabiochem), hexafluorophosphate benzotriazole tetramethyl uronium (Anaspec), and 1-hydroxybenzotriazole hydrate (Anaspec), and finally 10 equiv. DIPEA (Sigma-Aldrich) at 75 °C for 7 min. This procedure was used to couple all amino acids except cysteine, histidine, and arginine, which were coupled at 50 °C to limit racemization, and the coupling reaction was performed twice for arginine. The N-terminus of **1** used in CD experiments was not acetylated to best mimic the sample expressed recombinantly and used in NMR studies. **1** was cleaved from the resin and purified via reverse phase high performance liquid chromatography (RP-HPLC) as previously described^13^. The purity and observed mass of material isolated from pooled fractions was examined on an Agilent 6530 QTOF LCMS. Fractions were combined, lyophilized, and resuspended as previously described^13^.

### Expression and purification of ZF5.3 (3)

The gene fragments encoding ZF5.3 (**3**) (**Oligo 1** and **Oligo 2**) were cloned into a pET27b+ plasmid, linearized with **Primer 1** and **Primer 2,** using Gibson assembly according to manufacturer’s guidelines. The resultant plasmid His6-SUMO-ZF5.3 was verified by Sanger sequencing. Relevant DNA and protein sequences are listed in **Table S2**. His_6_-SUMO-ZF5.3 was transformed into *Escherichia coli* BL21DE3 Gold (Agilent) and selected on a kanamycin Luria Broth (LB) agar plate. Briefly, a single colony harboring His_6_-SUMO-ZF5.3 was picked and grown in 5 mL LB supplemented with 150 mg/L kanamycin at 37 °C for 6-8 h. Next, 1 mL of this culture was used to inoculate 500 mL of minimal media (50 mM Na_2_HPO_4_, 20 mM KH_2_PO_4_, 9 mM NaCl, 4 g/L glucose, 1 g/L NH_4_Cl, 0.1 mM CaCl_2_, 2 mM MgSO_4_, 10 mg/L thiamine, 10 mg/L biotin, and 150 mg/L kanamycin pH 7.4) and allowed to grow overnight with shaking at 37 °C. The next day, 950 mL of minimal media were inoculated with 50 mL of the overnight culture and allowed to grow at 37 °C to an OD_600_= 0.6-0.8 at which expression was induced with 1 mM isopropyl β-D-thiogalactopyranoside and transferred to 18.5 °C for 12-16 h. All subsequent steps were performed at 4 °C unless otherwise noted. Cells from the expression cultures described above were harvested by centrifugation at 4,300 rpm for 30 min, resuspended in 25 mL **Lysis Buffer** containing 2 Roche cOmplete mini protease inhibitor tablets (Sigma-Aldrich) and lysed via high pressure homogenization on a EmulsiFlex-C3 homogenizer. The cell lysate was clarified by centrifugation at 10,000 rpm for 45 min. The soluble supernatant was incubated with TALON® affinity resin, pre-equilibrated with lysis buffer, for 1 h. The resin was washed thoroughly with 50 mL **Wash Buffer** and His_6_-SUMO-ZF5.3 was eluted with 20 mL **Elution Buffer**. All protein concentrations were determined using a commercially available Pierce 660 nm protein quantification kit. Pooled elution fractions were combined with 1/16 equivalents of SUMO protease and dialyzed in dialysis tubing with a molecular weight cut off (MWCO) of 1,000 Da (Spectrum) against **Dialysis Buffer** at room temperature for 2 h. All column purifications were performed by fast performance liquid chromatography on an ÄKTA Pure (Cytiva). The cleavage reaction was then directly loaded onto an 5 mL HiTrap™ SP HP cation exchange column (Cytiva) and purified over 20 column volumes with a linear gradient of 0 to 100% **Cation Exchange Buffer B**. Fractions corresponding to pure ZF5.3 were then treated with 2 equivalents of ZnCl_2_ at room temperature for 15 min and concentrated and buffer exchanged via spin centrifugation at 4,000 rpm 4 °C with spin filters of 2,000 Da MWCO (Vivaspin). For CD experiments, ZF5.3 was buffer exchanged into **Resuspension Buffer**.

### Preparation and characterization of ^15^N-ZF5.3 for Heteronuclear Quantum Coherence Spectroscopy (HSQC) experiments

^15^N-ZF5.3 was generated recombinantly as previously described, with the only difference being the use of ^15^NH_4_Cl (Cambridge Isotopes) in the minimal media formulation. For NMR experiments with ^15^N-ZF5.3, the protein was concentrated to 800 µM and buffer exchanged into **^15^N-^1^H HSQC Buffer**. The pH of these samples was measured and modulated in the same fashion as the CD experiments. The homogeneity of ^15^N-ZF5.3 was analyzed via analytical size-exclusion chromatography. The molecular weight was examined on an Agilent 6530 QTOF LCMS.

### Preparation and characterization of ^13^C & ^15^N-ZF5.3 for triple resonance backbone and side-chain assignment experiments

^13^C & ^15^N-ZF5.3 was generated recombinantly as previously described, with the only difference being the use of U-^13^C glucose and ^15^NH_4_Cl (Cambridge Isotopes). For NMR experiments with ^13^C & ^15^N-ZF5.3, the protein was concentrated to 800 µM and buffer exchanged into **Structure Determination Buffer**. The homogeneity of ^13^C & ^15^N-ZF5.3 was analyzed via analytical size-exclusion chromatography. The molecular weight was examined on an Agilent 6530 QTOF LCMS.

### Synthesis of Rho-DBCO and Rho-ZF5.3 (1^R^)

Rhodamine labeling of ZF5.3 (synthesized on Rink Amide resin (100-200 mesh) Novabiochem®) was achieved by the incorporation of (S)-2-(Fmoc-amino)-6-azidohexanoic acid at the peptide N-terminus followed by acetylation to generate Ac-Lys(N_3_)-ZF5.3. The peptide was cleaved from the resin with 82.7% TFA, 5.1% phenol, 5.1% H_2_O, 5.1% thioanisole, 1% DTT (w/v), and 1% TIPS^13^. Following cleavage and purification by RP-HPLC, Ac-Lys(N_3_)-ZF5.3 was reacted with Lissamine Rhodamine B ethylenediamine functionalized with a DBCO (Rho-DBCO). Rho-DBCO was generated by reacting Lissamine Rhodamine B ethylenediamine (#L2424) with 10 molar equivalents of DBCO-NHS ester (#CCT-1491) in DMSO at RT for 1 h. HPLC-purified Rho-DBCO was reacted with Ac-Lys(N_3_)-ZF5.3 under the same conditions stated above. Final Rho-ZF5.3 was obtained following RP-HPLC and lyophilized and reconstituted as previously described^13^.

### Synthesis of BBA5.3 (2)

**2** was synthesized in a similar fashion to **1** with the following minor modifications. Rink amide was used and 0.1M Hydroxybenzotriazole (Anaspec) was included in the deprotection solution to limit aspartimide formation. In addition, amino acids following D-Pro were double coupled. Acetylation of the N-terminus was achieved as previously described^13^. Cleavage and purification were performed under the same conditions as **1**. Rhodamine labeling of BBA5.3 was achieved by the incorporation of an N-terminal Boc-Lys(Fmoc)-OH monomer which was selectively deprotected with 20% 4-methyl piperidine and reacted with Lissamine Rhodamine B Sulfonyl Chloride as previously described^13^.

## CD

### Circular dichroism

All circular dichroism (CD) data were collected on an AVIV Biomedical, Inc. (Lakewood, NJ) Circular Dichroism Spectrometer Model 410 or a JASCO J-1500 Circular Dichroism Spectropolarimeter. Wavelength-dependent CD spectra were recorded between 200 and 300 nm, measuring every 1 nm with an average sampling time of 5 seconds in a 1 mm cuvette. All measurements were normalized to a baseline buffer-only spectrum (Starna). For wavelength spectra acquired across multiple pHs, the pH of the sample was adjusted with 0.1M HCl or 0.1M KOH and measured on a pH sensor InLab® Micro (Mettler-Toledo) before and after the spectra were recorded. All pH values were within ± 0.1 pH units. For temperature melt experiments, ellipticity was measured at 208 nm from 2 °C to 95 °C. The average raw ellipticity of three technical replicates was used to determine the mean residue ellipticity (MRE) reported using the equation (1).

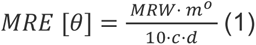

where *θ* = mean residue ellipticity in 10^3^*deg*cm^2^*dmol^-1^, MRW = mean residue weight (molecular weight divided by the number of backbone amides), m° = raw ellipticity, c = protein concentration in mg/mL, and d is cuvette path length in cm.

### Circular dichroism of zinc-free ZF5.3 (1)

Lyophilized ZF5.3 (**1**) was used in titration experiments without Zn(II) present to eliminate any complications associated with the chelation of Zn(II) in the ZF5.3 (**3**) sample expressed recombinantly. Wavelength experiments were performed under the exact same conditions as described above however in the absence of Zn(II).

## NMR

### ^15^N-^1^H HSQC experiments on ^15^N-ZF5.3

^15^N-^1^H HSQC spectra were measured on a Bruker 900 MHz Avance II spectrometer equipped with TCI cryoprobe.

### 15N-1H HSQC with added delay experiments on ^15^N-ZF5.3

The chemical shifts of ^1^H and ^15^N atoms in His side chains of ZF5.3 were determined on the same ^15^N-ZF5.3 samples used in the 15N-1H HSQC experiments stated above however with the addition of a 22 ms delay (1/(2JNH)) to refocus single-bond correlations^18^.

### ZF5.3 pH 5.5 structure determination

Two- and three-dimensional (2D/3D) spectra for backbone (N^15^HSQC, CBCAcoNH, CBCANH) and side-chain assignment (^13^C-CTHSQC-aliphatic, HBHAcoNH, hCccoNH, ^13^C-HSQC-aromatic hbCBcgcdHD-aromatic, hbCBcgcdceHE-aromatic) were recorded at 25 °C on a 600 MHz Bruker Avance NEO spectrometer equipped with an actively shielded Z-gradient 5 mm TCI Cryoprobe using programs from the TopSpin 4.0.6 pulse program library. A 3D hCCH-TOCSY experiment (T_m_ = 36 ms) and a 120 ms mixing-time simultaneous evolution 3D ^13^C-/^15^N-NOESY-HSQC experiment was recorded to measure distance restraints on on a 800 MHz Bruker Avance NEO spectrometer equipped with an actively shielded Z-gradient 5 mm TCI Cryoprobe at 25 °C, using programs from the TopSpin 4.0.8 pulse program library (TopSpin 4.0.6). Spectra were processed in TopSpin 4.3.0 and inspected using CcpNmr (ver. 3.2.0) Assign. Proton chemical shifts were referenced externally to a DSS standard while ^13^C and ^15^N chemical shifts were referenced indirectly to this value^52^. All experiments were carried out at 25 °C and the temperature calibrated in the standard way^53^. Structure determination utilized an AI-assisted automatic NMR structure determination pipe-line ARTINA^23^ implemented on the NMRtist webserver (https://doi.org/10.1093/bioinformatics/btad066), which utilizes a trained deep neural network to identify signals in NMR spectra, resonance assignments using FLYA^25^ and automatic nOe assignments and automatic structure determination using CYANA^27^. The final NMR ensemble was calculated based on 392 nuclear Overhauser effect restraints (nOe; 90 long-range nOes (|*i-j*| > 5), 81 medium-range nOes (1 < |*i-j*| < 4), 100 sequential nOes (|i-j| = 1), and 121 intra-residue nOes (|i-j| < 1) (**Extended Data Table 2**) and the initial set of twenty lowest-energy ZF5.3 conformers superposes with an average pairwise backbone root mean square deviation (RMSD) of (for residues 1 to 27) of 0.8 Å and 1.34 Å for backbone and heavy atoms, respectively (**Fig. 3a**)^23^.

### ZF5.3 pH 5.5 structure refinement

The final refinement involved a restrained MD refinement and then a final step with explicit water using the NIH-XPLOR (3.5) scripts refine.py and wrefine.py modified to include patches for the Zn-finger (zn-finger-allhdg5.3.par;bond lengths and angles were defined to be 2.3 Å and 109° Zn-Sγ-Cβ for Cys3 and Cys6 and 2.3 Å and 109° Zn-Nε-Cδ for His19 and His23). Two-hundred structures were calculated for each step and the final ensemble contained the 20 lowest energy structures. The final structures were validated using the wwPDB validation server (https://validate-rcsb-east.wwpdb.org/validservice/) and the Protein Structure Validation Suite (PSVS). Ramachandran statistics among the top 20 lowest-energy structures are 95.2% for most favored regions, 4.8% for additionally allowed regions and 0% for generously allowed regions and disallowed regions. The structure calculation statistics for the 20 lowest-energy structures are in **Extended Data Table 2**. We deposited this ensemble (All heavy atom RMSD = 1.1 Å, for the ordered residues: 2-24) and underlying NMR data to the PDB and BMRB with accession codes 9AZI and 31151, respectively.

## Cell studies

### Delivery of Rho-ZF5.3 (1^R^) and Rho-BBA5.3 (2^R^) to Saos-2 cells

Delivery of rhodamine-labeled peptides was performed as previously described^13^. Briefly, for delivery into Saos-2 cells, 0.2 x 10^6^ Saos-2 cells were plated in a 6-well plate in clear McCoy’s 5A media (+15% FBS) the day before experiments. The next day, 1 mL of Rho-labeled protein was added to each well at final concentrations of 1-3 µM diluted with McCoy’s 5A medium (-FBS, -phenol red). Incubations were allowed to go for 1 h at 37 °C, 5% CO_2_. In the final 5 min, a final concentration of 300 nM Hoechst 33342 nuclear stain was added to each well. After protein delivery, cells were washed 3 times with 2 mL of DBPS per wash and treated with 0.5 mL trypsin (TrypLE, -phenol red) to remove any surface bound peptide and lift cells for flow cytometry and microscopy analysis. Cells were buffer exchanged into either DPBS or DMEM (-FBS) for flow cytometry and microscopy, respectively.

### Flow cytometry and fluorescence correlation spectroscopy (FCS)

Flow cytometry analysis was performed at room temperature on an Attune NxT flow cytometer. Hoechst 33342 was excited using the violet laser at 405 nm, and the emission was recorded between 390 and 490 nm. Rhodamine was excited using the yellow laser at 561 nm, and the emission was recorded between 570 and 602 nm. The median fluorescence intensity (MFI) for the Rhodamine channel was reported for 10,000 gated cells, with at least 2 biological replicates per condition. Confocal microscopy and FCS were performed as previously described^10^. Briefly, all experiments were performed at 37 °C and 5% CO_2_ on a STELLARIS 8 microscope (Leica Microsystems) equipped with a HC Plan-Apo 63x/1.4NA water immersion objective (used for FCS measurements). A confocal microscopy image was used to position the microscope laser for FCS in discrete locations within the cytosol of 30-40 cells per well. For each point, ten 5-second autocorrelation traces were measured. Autocorrelation curves were fitted to derive concentration values as previously described using a custom MATLAB® script available from GitHub (https://github.com/schepartzlab/FCS)^13^. A minimum of two biological replicates were performed for each condition. After data filtering and fitting, there were at least 15 FCS points used to determine mean cytosolic concentration for each condition.

## Liposomes

### Preparation and characterization of large unilamellar vesicles (LUVs)

Lipids for preparation of LUVs first were dissolved in chloroform and mixed to the molar ratios described in **Table S2** (10 µmol total lipid). Rhodamine-labeled LUVs (Rho-LUVs) were prepared by addition of 1 nmol 1,2-dioleoyl-sn-glycero-3-phosphoethanolamine-N-(lissamine rhodamine B sulfonyl (Avanti #810150) to the lipid mixture. Chloroform was evaporated under N_2_ to produce a thin lipid film. The resultant film was hydrated with 1 mL buffer (20 mM Tris-HCl, 150 mM KCl, pH 7.5 or pH 4.5) for 30 min, then vortexed for 1 min to fully redissolve. Next, the lipid mixture was freeze-thawed for 3 cycles followed by 11 passes through a 200 nm polycarbonate (Avanti Polar Lipids #610006) using an Avanti Mini Extruder (Avanti Polar Lipids #610023). LUV diameter was confirmed via dynamic light scattering and stored at 4 °C for use within 72 hours of preparation. DLS measurement was performed on a Malvern Instruments Zetasizer Nano ZS. LUV capsid samples were prepared at a 1 mM lipid concentration in buffer (20 mM Tris-HCl, 150 mM KCl, pH 7.5) and passed through a 0.2 μM spin filter prior to each run. Size determination was performed in triplicate at 25 °C following a 5-min temperature equilibration.

### Liposomal co-sedimentation assay

The liposomal co-sedimentation assay was adapted from a previously reported protocol with modifications for a fluorescence-based readout.^42^ To 0.2 mL ultracentrifuge tubes (Beckman Coulter #343775) 1 µM mini-protein **1^R^**or **2^R^** and 5 mM **ILV, BMP+LE/LY, BMP-LE/LY,** or **PM** LUVs was added in **Resuspension Buffer** (adjusted to pH 7.5 or 4.5). Samples were incubated for 30 min at 37 ℃, then LUVs (along with bound mini-protein) were pelleted by ultracentrifugation (TL-100; Beckman Coulter, TLA-100 rotor) at 100,000g for 30 min. Supernatant (containing unbound mini-protein) was transferred to a clear bottom 96-well plate (Corning #3904). The remaining pellet was resuspended in an equal volume of **Resuspension Buffer** and transferred to the 96-well plate. The fluorescence was measured using an Agilent BioTek Synergy H1 Microplate Reader (excitation: 545 nm / emission: 586 nm). The fluorescence of mini-protein pelleted in the absence of LUVs was subtracted from each sample as background. Fractional binding of mini-proteins was determined using equation 2 and each sample was prepared in triplicate.

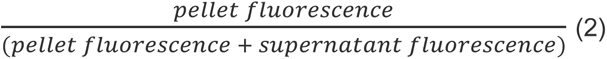

### Mini-protein-LUV binding FCS

FCS for mini-protein-LUV binding determination was performed as described in **Methods in support of Figure 5 and Extended Data Figure 8** with the following modifications. First, single-component *in vitro* FCS was performed to determine the mini-protein and LUV diffusion times. Mini-proteins **1^R^** and **2^R^** were diluted to 100 nM in **Resuspension Buffer** and **Rho-ILV, Rho-BMP+LE/LY, Rho-BMP-LE/LY,** and **Rho-PM** LUVs were diluted to 500 µM lipid in buffer (20 mM Tris-HCl, 150 mM KCl, pH 7.5). Ten consecutive, five-second FCS measurements were recorded for each sample. Raw autocorrelation and countrate data were averaged and fitted to a 3D diffusion equation (eq 3) to obtain the *in vitro* diffusion time of each mini-protein or LUV.

For two-component binding FCS (**Fig. 6c**) mini proteins **1^R^**or **2^R^** were mixed with **ILV, BMP+LE/LY, BMP-LE/LY,** or **PM** LUVs in **Resuspension Buffer** (100 nM mini-protein, 500 µM lipid) and incubated for 30 min at 37 ℃. Ten consecutive, 30-second FCS measurements were recorded for each sample. Each curve was fitted individually to a 3D two-component diffusion equation (eq 4) with the 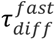 component fixed to the measured *in vitro* values for each mini-protein (0.63 ms or 0.54 ms for **1^R^**or **2^R^**, respectively) and 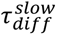 was constrained between 1-10 ms to encompass the range of diffusion times exhibited by the LUV populations. From each fit, (1-F_fast_) represents the fraction of mini-proteins in the focal volume bound to an LUV. Equilibrium binding curves (**Fig. 6d-e**) were produced using the same protocol with varying concentrations of mini-protein (60 pM-500 nM) with lipid concentration held constant (500 µM).

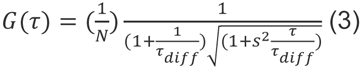

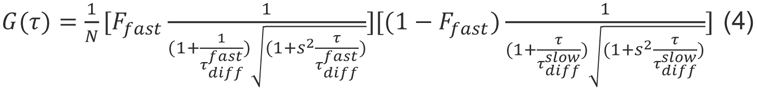

## Acknowledgements

This work was supported by the National Science Foundation (CHE-2203903 to A.S.), the National Institutes of Health (GM134963 to A.S. and 1S10OD023455-01A1 (UCSF NMR core facility)), and a PBBR TMC award (UCSF NMR core facility). This material is based upon work supported by the National Science Foundation Graduate Research Fellowship Program (NSF grant number: DGE 2146752 awarded to J.G.) and the National Science Foundation MPS-Ascend Postdoctoral Research Fellowship (NSF grant number: CHE-2213241 awarded to D.B.). Any opinions, findings, and conclusions or recommendations expressed in this material are those of the author(s) and do not necessarily reflect the views of the National Science Foundation. We thank members of the Schepartz lab for fruitful discussions and Charles Schwieters for advice on Xplor calculations with Zn-ligands.

## Author contributions

J.G., D.B., M.K., and A.S. designed the project. J.G. led NMR experiments and experiments involving BBA5.3. D.B. led experiments with liposomes. M.Z. designed and performed CD experiments. A.V. synthesized and delivered Rho-ZF5.3. M.K. oversaw all NMR experiments. J.G., D.B., M.K., and A.S. analyzed experiments and prepared the manuscript.

## Additional Information

### Extended Data

**Extended Data Figs. 1-8.**

**Extended Data Tables. 1-2.**

### Supplemental Data

**Supplemental Fig. 1.**

**Supplemental Data Tables. 1-5.**

---

**Extended Data Fig 1.**
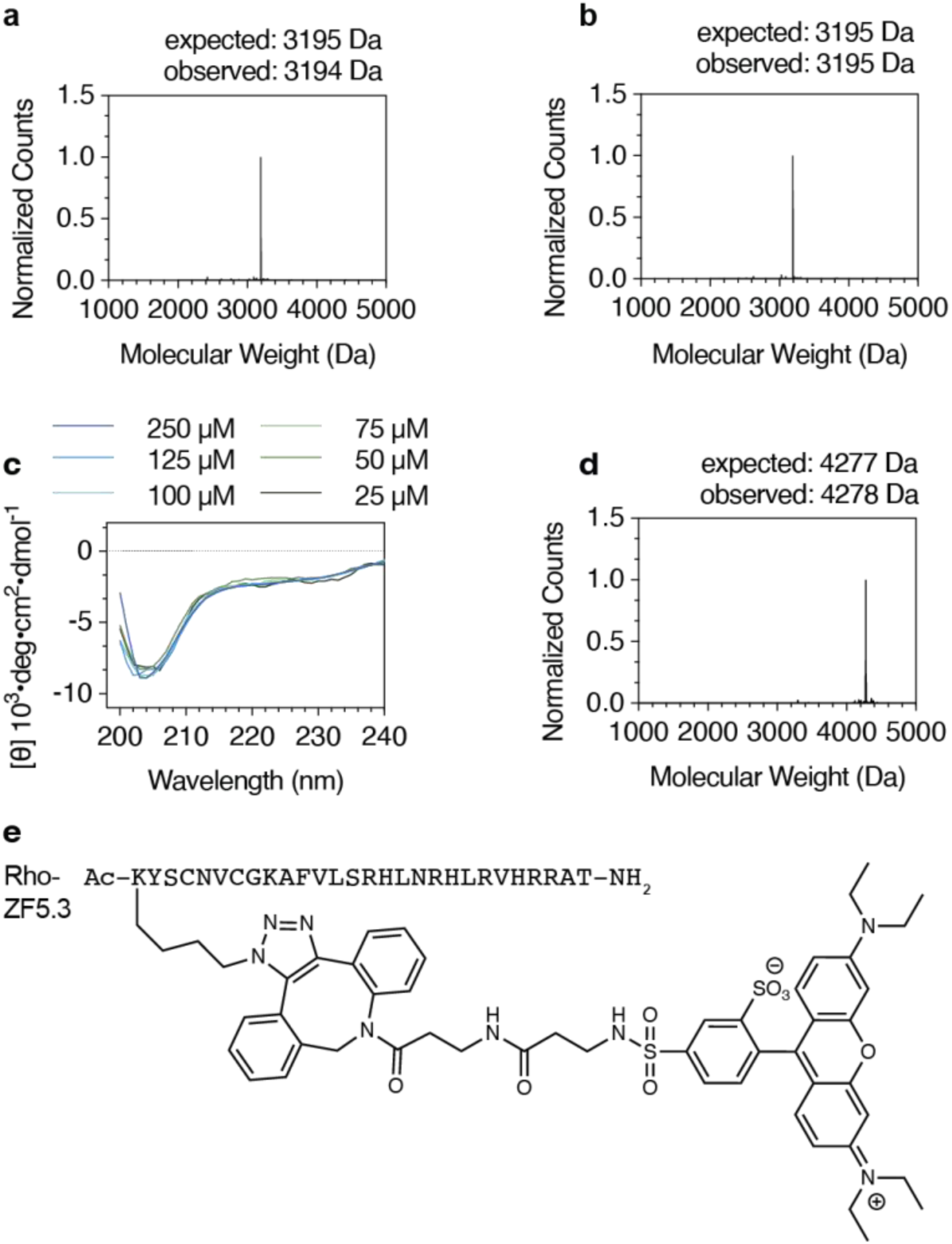
Additional data in support of. Figure 1. Deconvoluted mass spectra of ZF5.3 generated **a,** recombinantly or **b,** by solid phase synthesis as described in **Methods**. **c,** CD spectra of recombinant ZF5.3 in Reconstitution Buffer at pH 7.5 (20 mM Tris-HCl, 150 mM KCl, 1 mM TCEP, and 100 µM ZnCl_2_) at concentrations between 25 µM and 250 µM. **d,** Deconvoluted mass spectrum of Rho-ZF5.3 (pink, used for confocal microscopy, flow cytometry, and FCS experiments) prepared using solid-phase peptide synthesis as described in **Methods**. **e,** Sequence of Rho-ZF5.3.

**Extended Data Fig 2.**
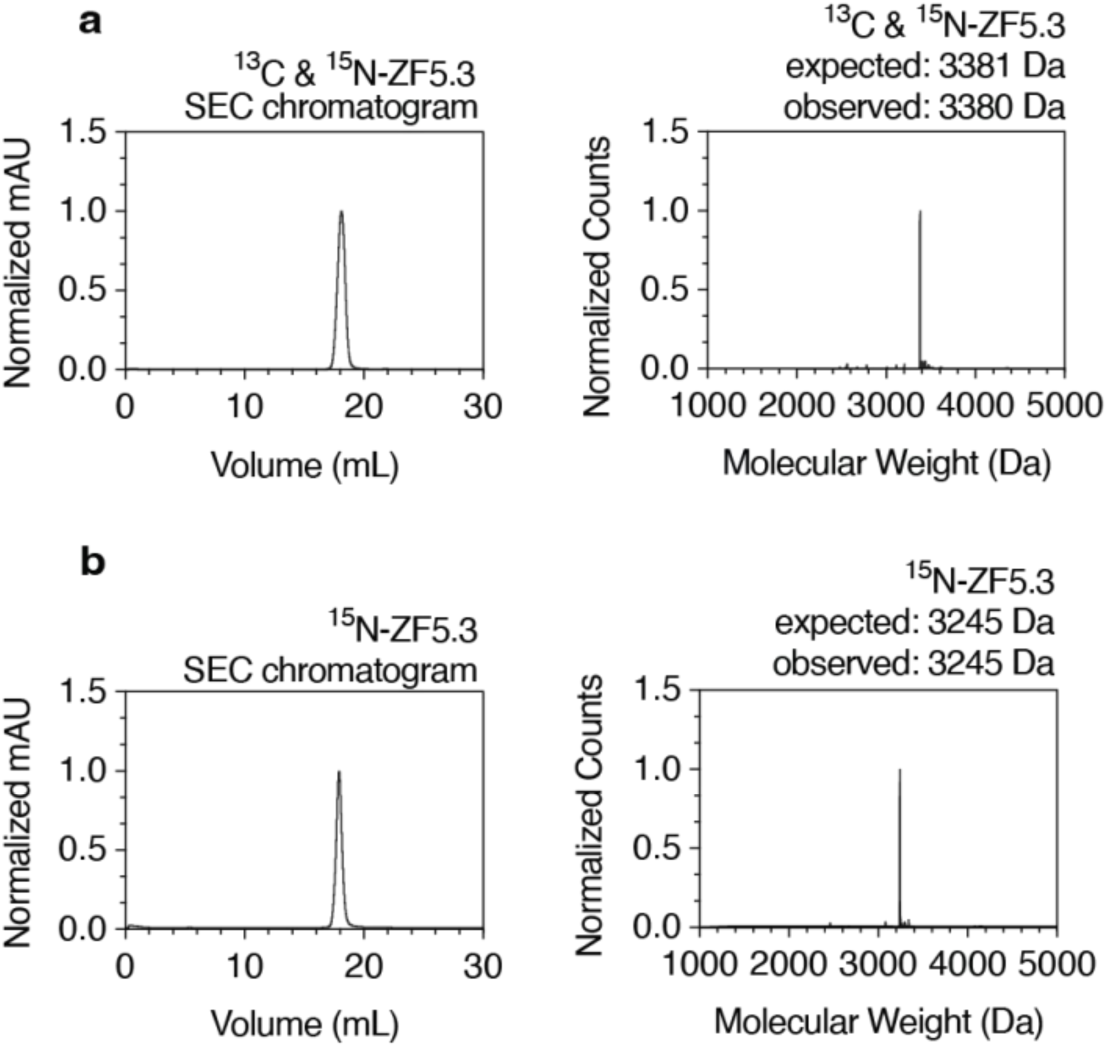
Characterization of recombinantly expressed and purified samples of ZF5.3 that were uniformly labeled with either **a,** ^15^N and ^13^C or **b,** ^15^N. Shown are analytical size-exclusion chromatograms (SEC) of each ZF5.3 sample following immobilized metal ion affinity and cation exchange chromatography alongside the deconvoluted mass spectrum. Additional experimental details are available in **Methods.**

**Extended Data Fig 3.**
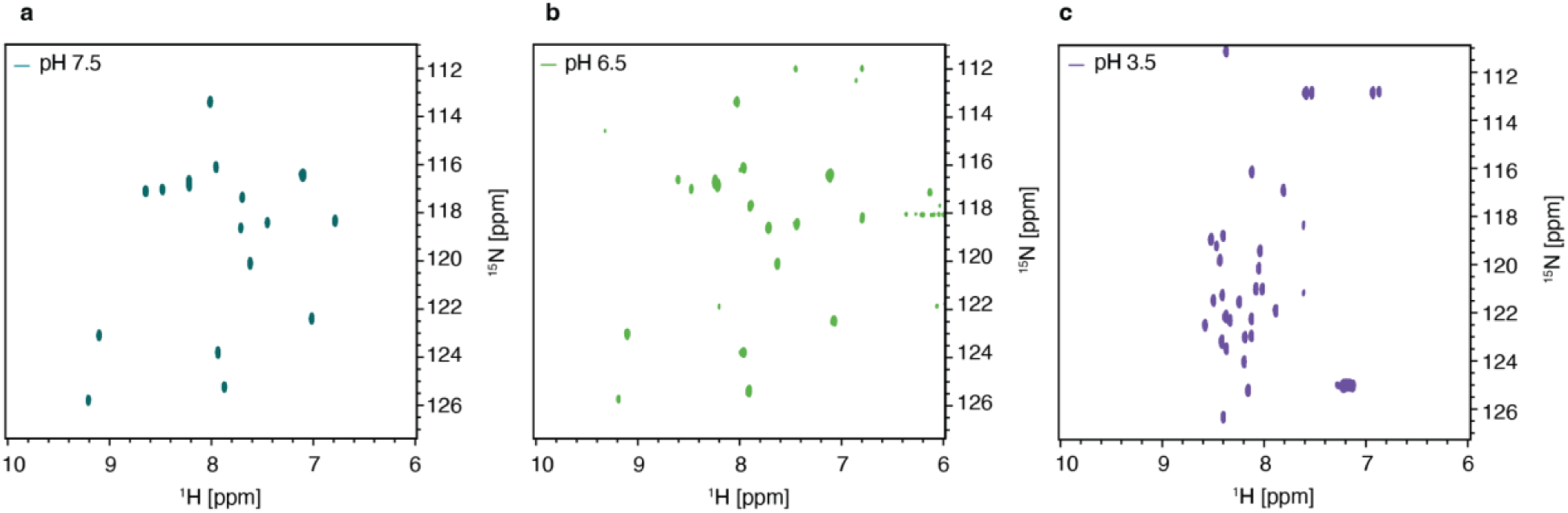
Additional data in support of. Figure 2**. a-c,**^15^N-^1^H heteronuclear quantum coherence (HSQC) spectroscopy at pH 7.5, 6.5, and 3.5. ^15^N-^1^H HSQC of ZF5.3 (800 µM) was acquired in a 20 mM citrate-phosphate buffer containing 10% D_2_O, 100 mM NaCl, 2 mM TCEP and 1.6 mM ZnCl_2_ at pH 7.5, 6.5, and 3.5. Additional experimental details are available in **Methods.**

**Extended Data Fig 4.**
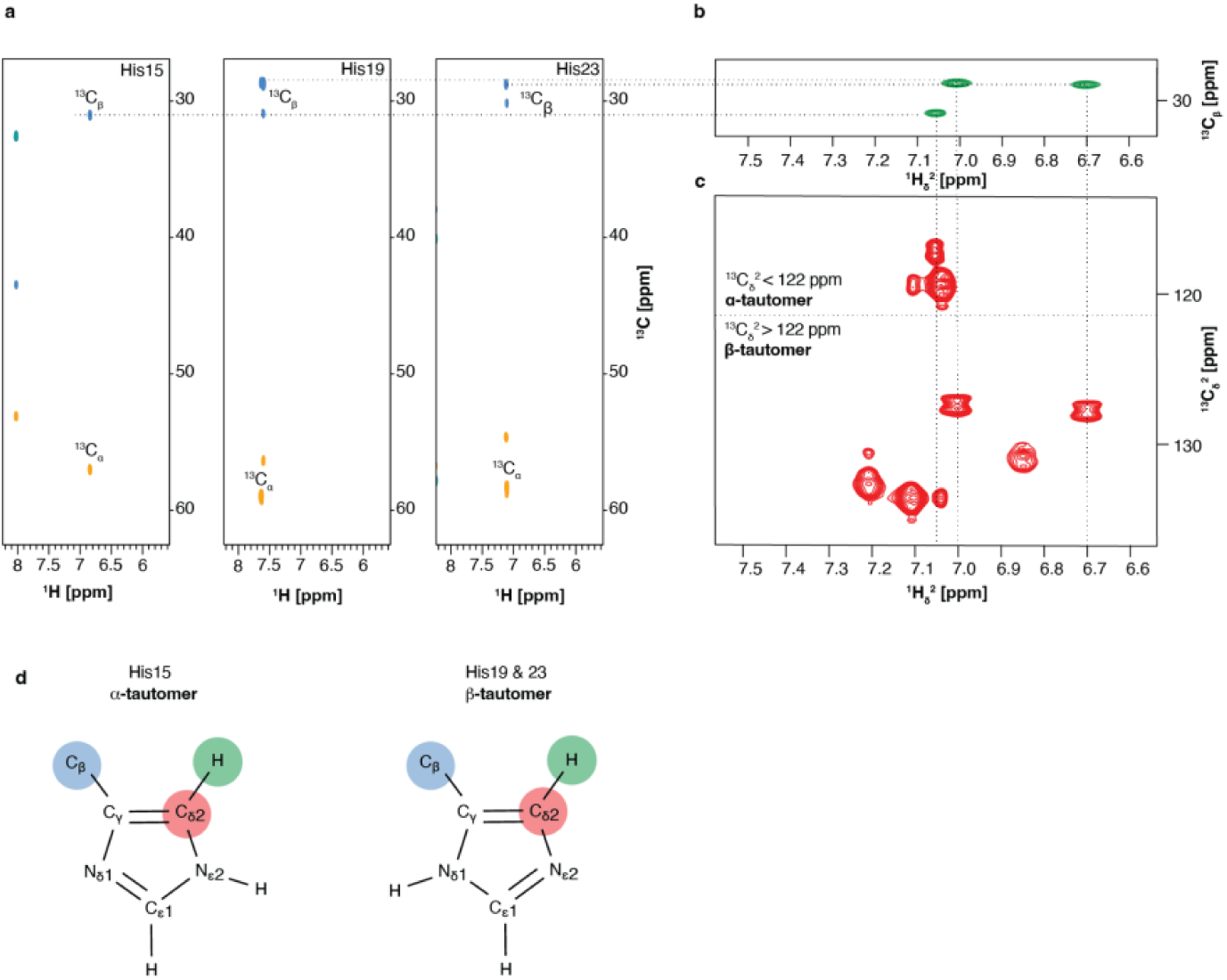
Assignment of His tautomers in ZF5.3 at pH 5.5. **a,** Results of hbCBcgcdHD experiments that provide the ^13^C⍺ and ^13^Cꞵ chemical shifts of each of the three His residues in ZF5.3, His15, His19, and His23. **b,** Results of a 2D CBHD experiment that correlates the chemical shift of each ^13^Cꞵ with the corresponding ^1^Hδ. **c,** ^13^C-CT HSQC provides the chemical shift of ^13^Cδ and ^1^Hδ. If the chemical shift of 13Cδ is less than 122 ppm the His residue is said to be in the ⍺-tautomeric state and if the chemical shift of 13Cδ is greater than 122 ppm it is in the ꞵ-tautomeric state. **d,** Assignment of His15 as the ⍺-tautomer and Zn(II) coordinating His19 and His23 as the ꞵ-tautomer.

**Extended Data Fig. 5.**
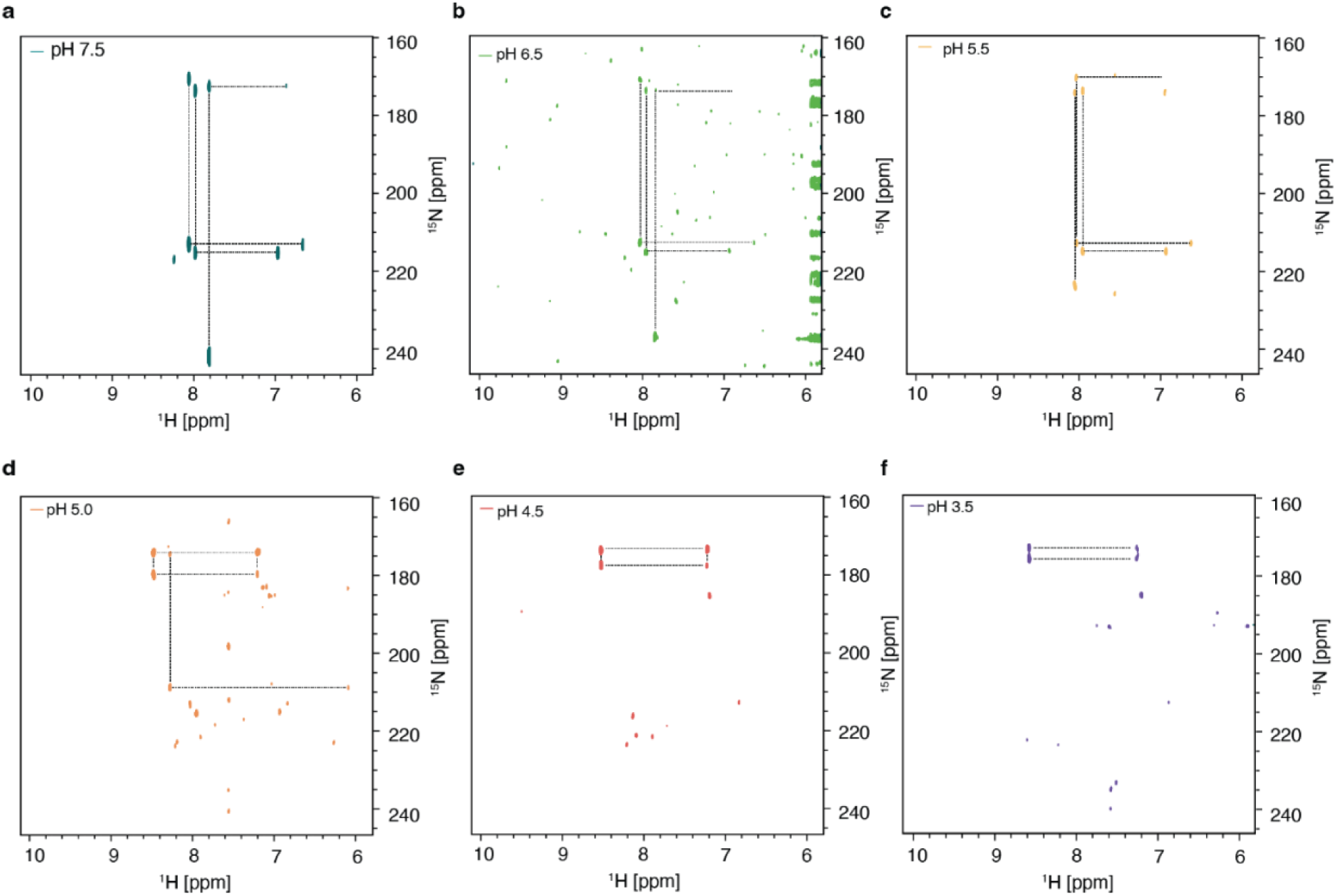
2D [^15^N-^1^H]-HMBC spectra provide evidence for His protonation at low pH. a-f, [^15^N-^1^H] heteronuclear multiple bond quantum coherence spectroscopy (HMBC) at pH 7.5, 6.5, 5.5, 5.0, 4.5, and 3.5 provides the chemical shifts of ^15^N and ^1^H atoms in His residues in ZF5.3.

**Extended Data Fig 6.**
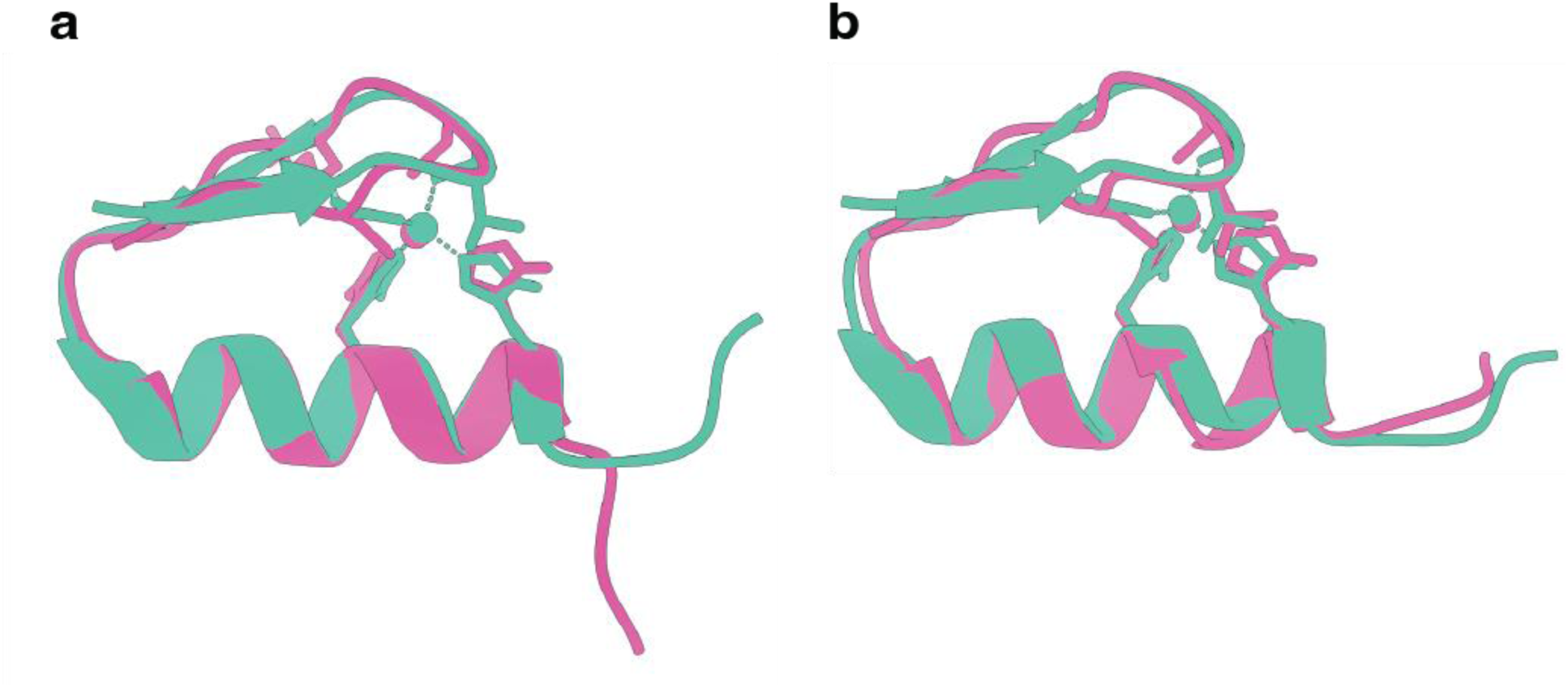
Alignment of ZF5.3 ensemble at pH 5.5 with ZF473. a, Overlay of a single ZF473 conformer (mint) with the ZF5.3 conformer (pink) that resulted in the lowest pruned atom pair RMSD value of 0.586 Å. **b,** Overlay of a single ZF473 conformer (mint) with ZF5.3 conformer (pink) with the lowest RMSD value across all 27 pairs of 0.812 Å.

**Extended Data Fig. 7.**
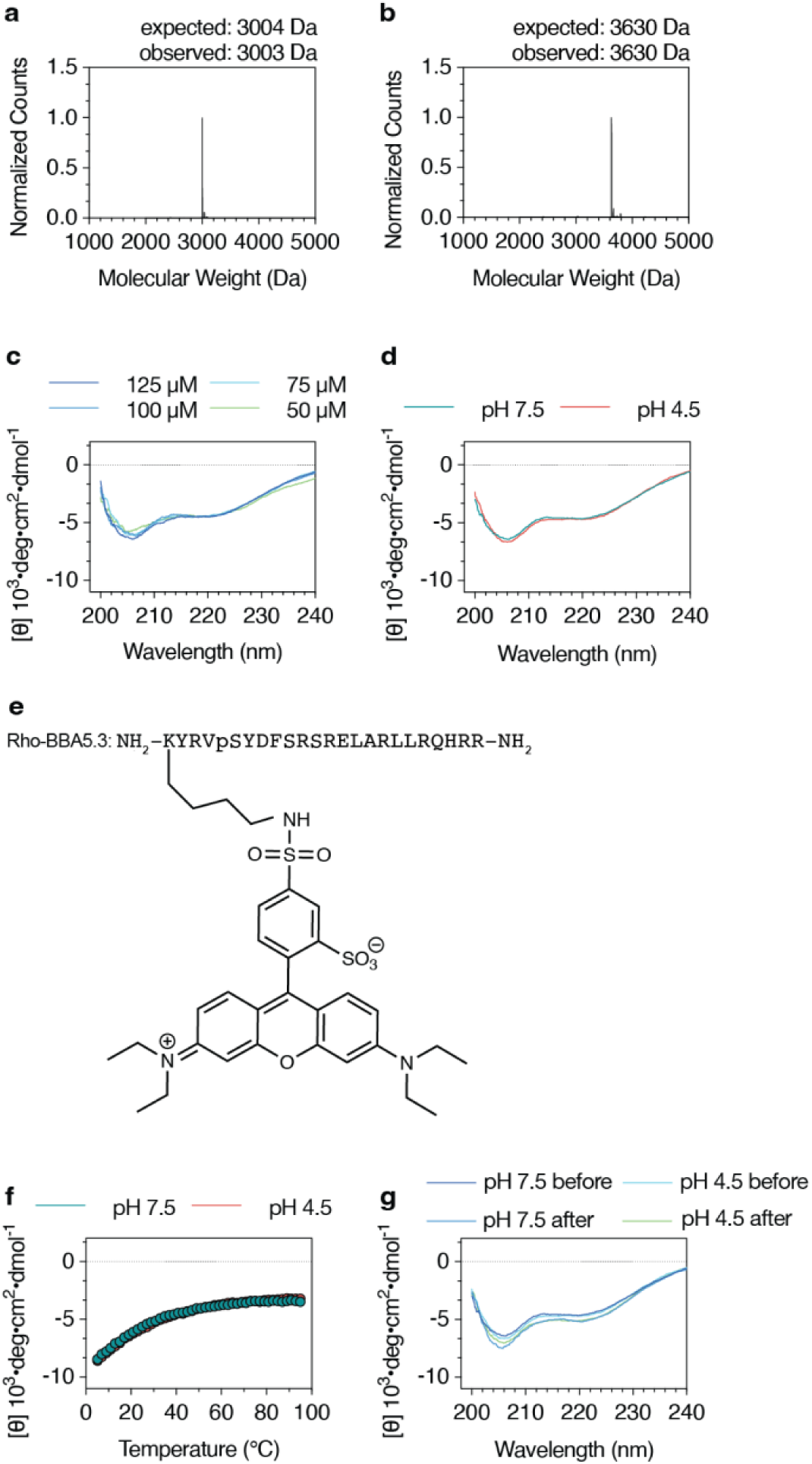
Additional data in support of Fig. 5. Deconvoluted mass spectra of HPLC-purified. **a,** BBA5.3 and **b,** Rho-BBA5.3. **c,** Overlay of the wavelength-dependent CD spectra of BBA5.3 at concentrations between 50 and 125 µM in a Resuspension Buffer without Zn(II) containing 20 mM Tris, 150 mM KCl, and 1 mM TCEP at 37 °C. **d**, Wavelength-dependent CD spectra of 125 µM (0.3 mg/mL) BBA5.3 at pH 7.5 and 4.5 in Resuspension Buffer without Zn(II) at 37 °C. **e**, Sequence of Rho-BBA5.3 prepared using solid phase synthesis as described in **Methods**. **f**, Plot of the mean residue ellipticity at 222 nm of 125 µM BBA5.3 in Resuspension Buffer without Zn(II) at pH 7.5 (teal) or pH 4.5 (red) at every 2 °C between 5 °C and 95 °C. **g,** Wavelength-dependent CD spectra of BBA5.3 (125 µM) before and after being heated from 5 to 95 °C.

**Extended Data 8.**
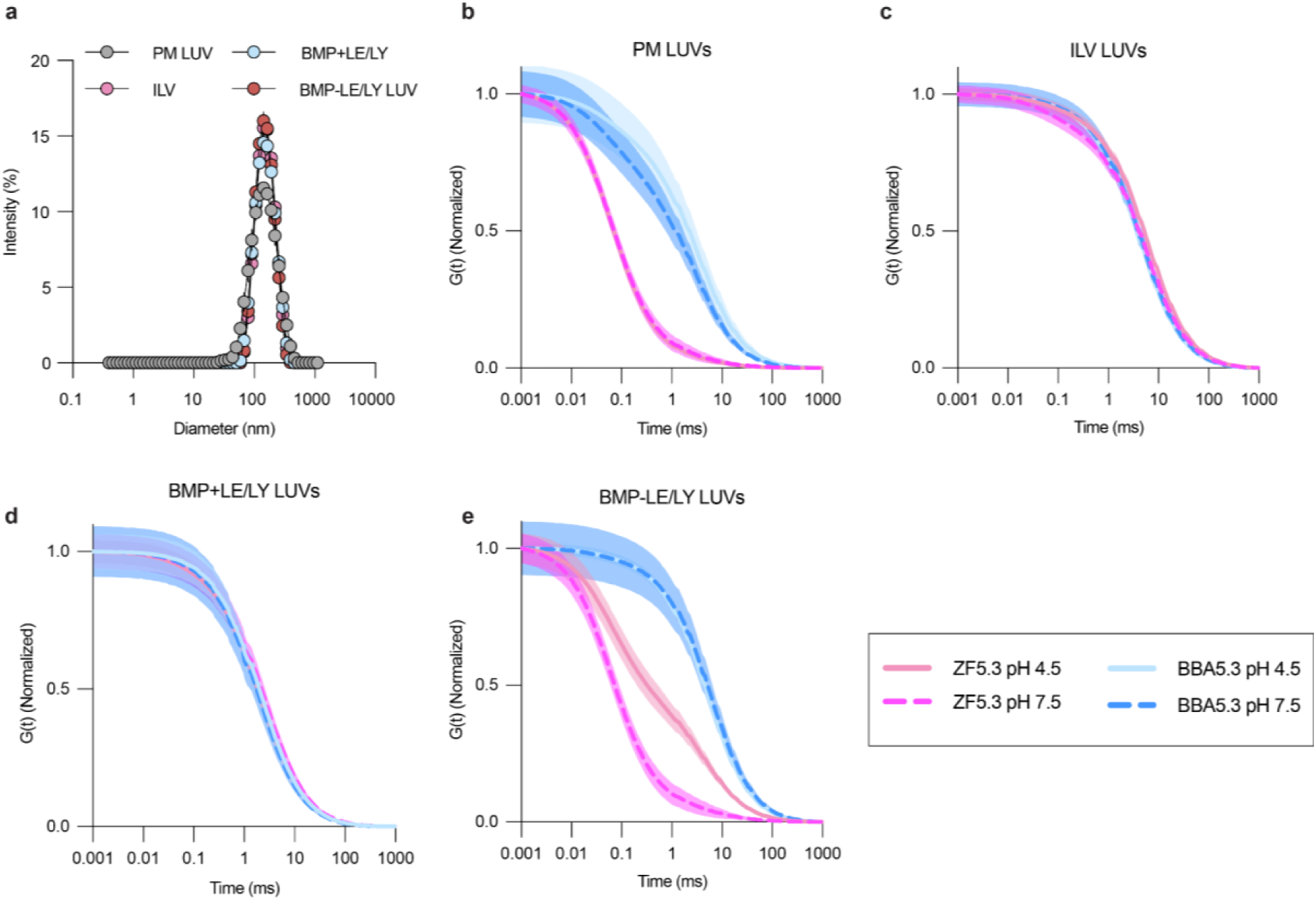
Data in support of Fig. 6. **a,** Diameter of PM, ILV BMP+LE/LY, and BMP-LE/LY LUVs determined by dynamic light scattering. Size distribution is reported as percent intensity (mean ± SEM), n=3. **b-e,** Normalized two-component fluorescence correlation spectroscopy autocorrelation curves (mean ± SEM) of mini-proteins Rho-ZF5.3 or Rho-BBA5.3 with PM, ILV,N BMP+LE/LY, and BMP-LE/LY LUVs at pH 4.5 or 7.5, n=10.

**Extended Data Table 1.**
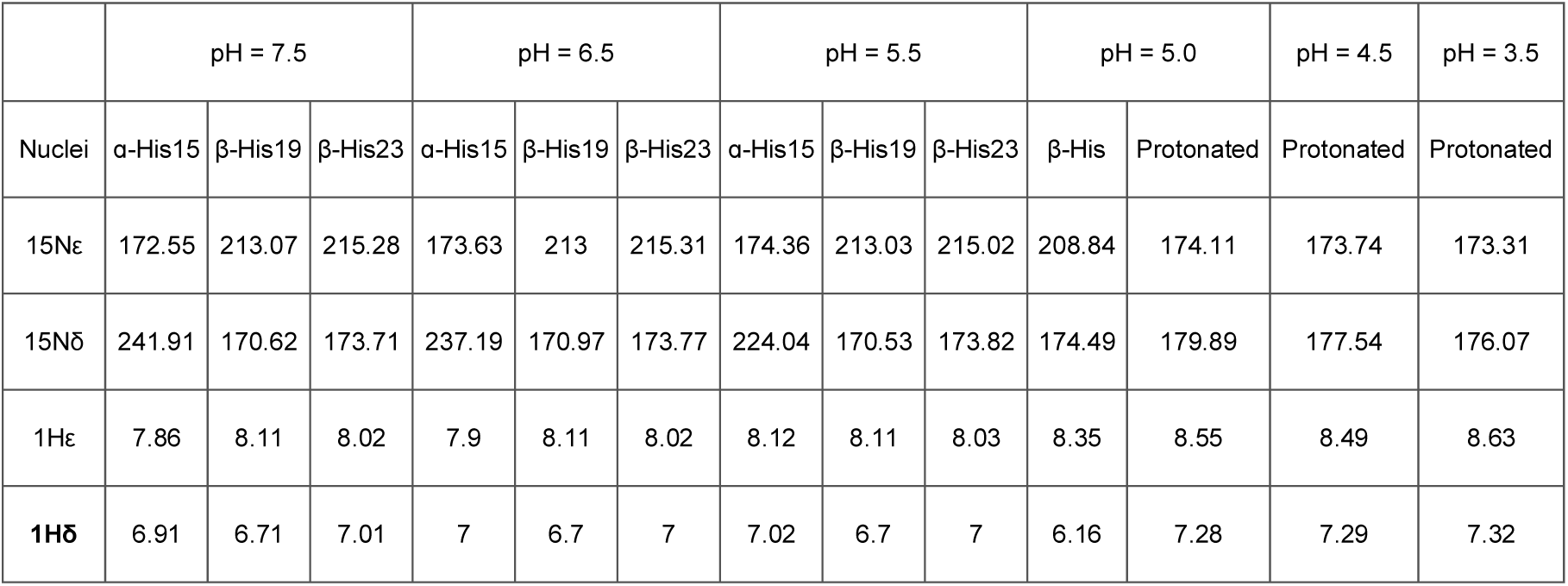
Tabulated values for ^1^Hδ, 1Hε, ^15^Nε, and ^15^Nδ for His residues in ZF5.3 at the indicated pH values between 7.5 to 3.5.

|  | pH = 7.5 |  |  | pH = 6.5 |  |  | pH = 5.5 |  |  | pH = 5.0 |  | pH = 4.5 | pH = 3.5 |
| --- | --- | --- | --- | --- | --- | --- | --- | --- | --- | --- | --- | --- | --- |
| Nuclei | α-His15 | β-His19 | β-His23 | α-His15 | β-His19 | β-His23 | α-His15 | β-His19 | β-His23 | β-His | Protonated | Protonated | Protonated |
| $^{15}\text{N}\epsilon$ | 172.55 | 213.07 | 215.28 | 173.63 | 213 | 215.31 | 174.36 | 213.03 | 215.02 | 208.84 | 174.11 | 173.74 | 173.31 |
| $^{15}\text{N}\delta$ | 241.91 | 170.62 | 173.71 | 237.19 | 170.97 | 173.77 | 224.04 | 170.53 | 173.82 | 174.49 | 179.89 | 177.54 | 176.07 |
| $^1\text{H}\epsilon$ | 7.86 | 8.11 | 8.02 | 7.9 | 8.11 | 8.02 | 8.12 | 8.11 | 8.03 | 8.35 | 8.55 | 8.49 | 8.63 |
| **$^1\text{H}\delta$** | 6.91 | 6.71 | 7.01 | 7 | 6.7 | 7 | 7.02 | 6.7 | 7 | 6.16 | 7.28 | 7.29 | 7.32 |

**Extended Data Table 2.** NMR and refinement statistics for ZF5.3 structure.

|  | ZF5.3 |
| --- | --- |
| **NMR distance and dihedral constraints** |  |
| Distance constraints |  |
| Total NOE | 392 |
| Intra-residue | 121 |
| Inter-residue | 271 |
| Sequential ($\vert i - j\vert = 1$) | 100 |
| Medium-range ($\vert i - j\vert < 4$) | 81 |
| Long-range ($\vert i - j\vert > 5$) | 90 |
| Intermolecular | 0 |
| Hydrogen bonds | 0 |
| Total dihedral angle restraints | 34 |
| **Structure statistics** |  |
| Violations (mean and s.d.) |  |
| Distance constraints (Å) | 0.0037 +/- 0.0213 |
| Dihedral angle constraints (°) | 0.1823 +/- 0.5775 |
| Max. dihedral angle violation (°) | 3.9186 |
| Max. distance constraint violation (Å) | 0.427 |
| Deviations from idealized geometry |  |
| Bond lengths (Å) | 0.0034 +/- 0.0053 |
| Bond angles (°) | 0.3943 +/- 0.4742 |
| Impropers (°) | 0.3485 +/- 0.7018 |
| Average pairwise r.m.s. deviation** (Å) |  |
| Heavy | 1.2 |
| Backbone | 1.8 |

**Supplemental Data Figure 1.**
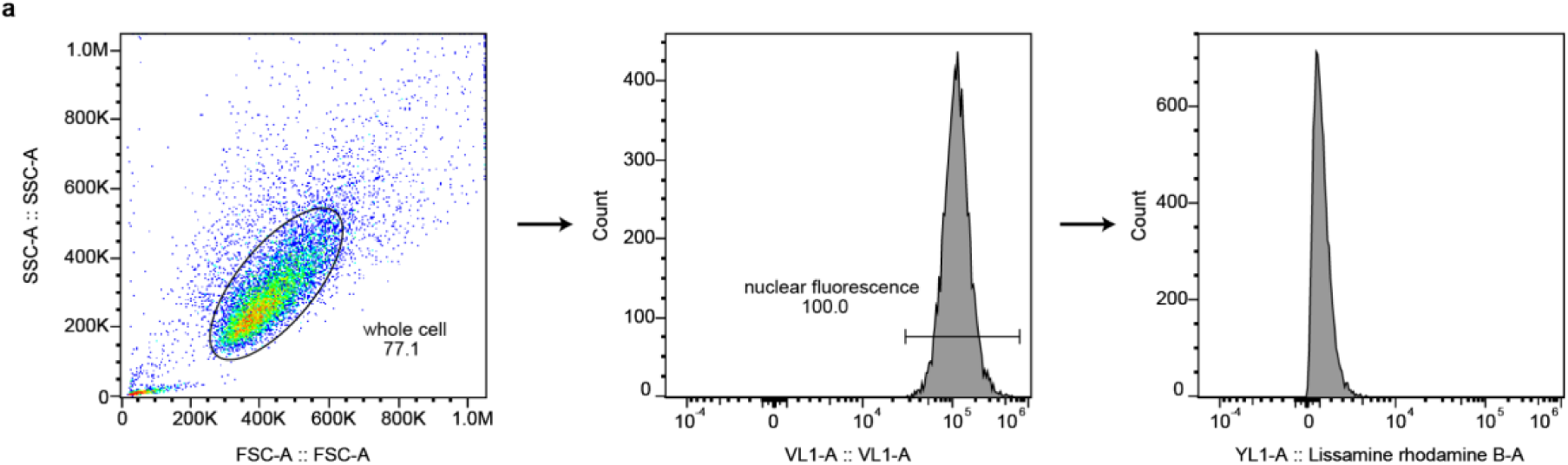
Flow cytometry gating and analysis for whole cells treated with Rho-BBA5.3 (1 hr at 37 °C, 5% CO2). Nuclei were identified by incubation with Hoechst 33342 for 5 min at the end of the incubation period. Cells were first gated based on FSC/SSC for the whole cell and selected for high nuclear fluorescence to obtain the Rhodamine fluorescence histogram.

**Table Supplemental Data Table 1 (S1).**
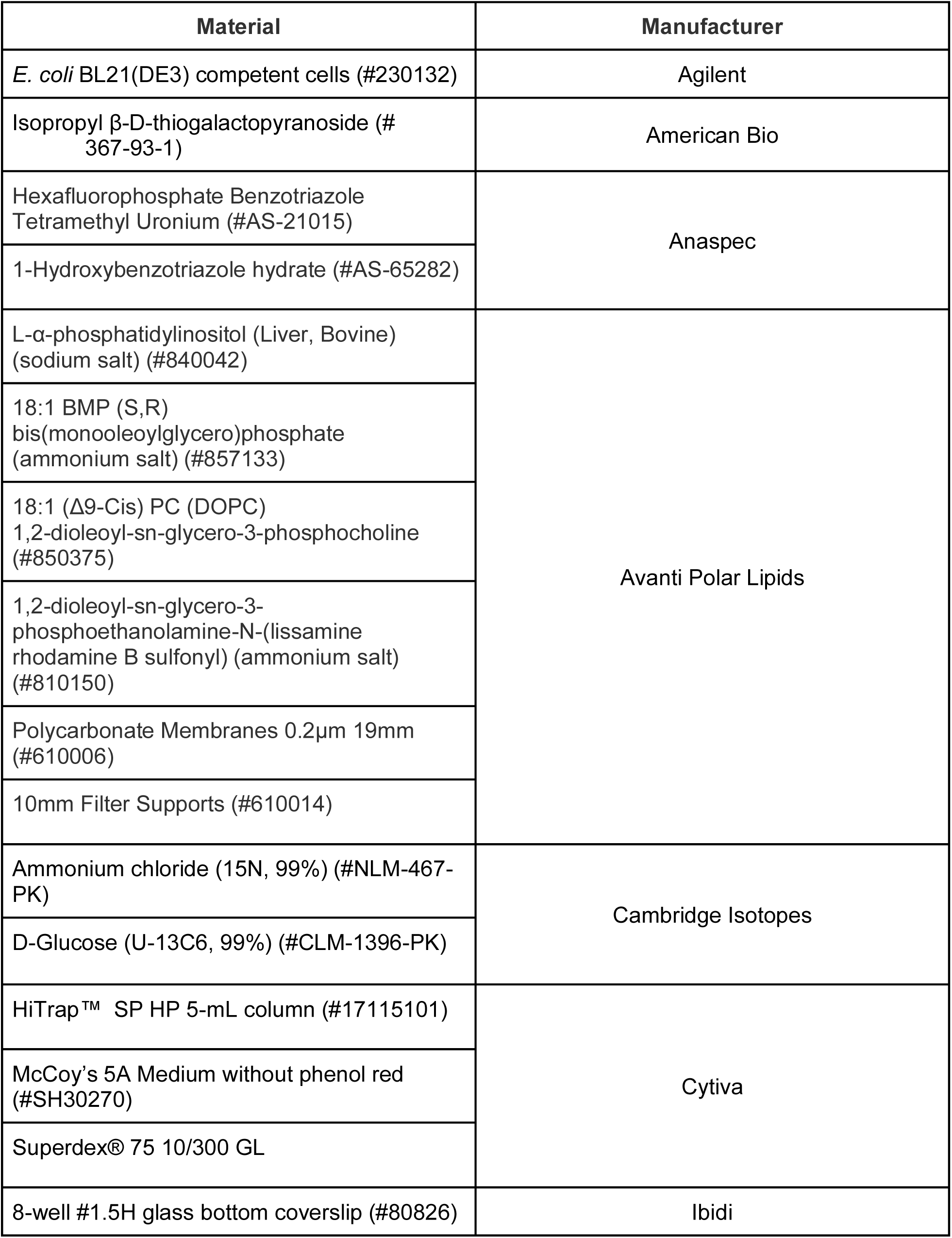
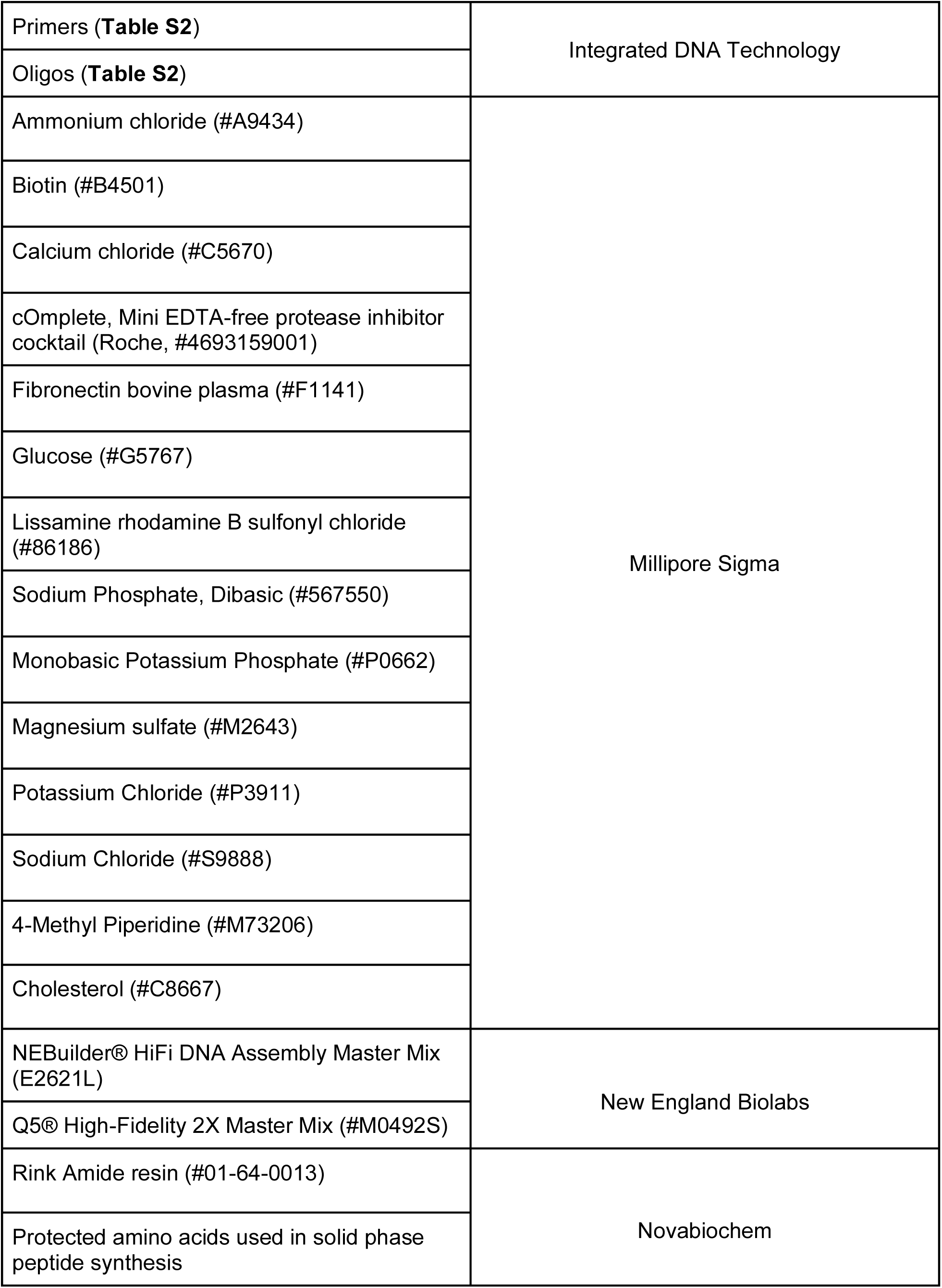
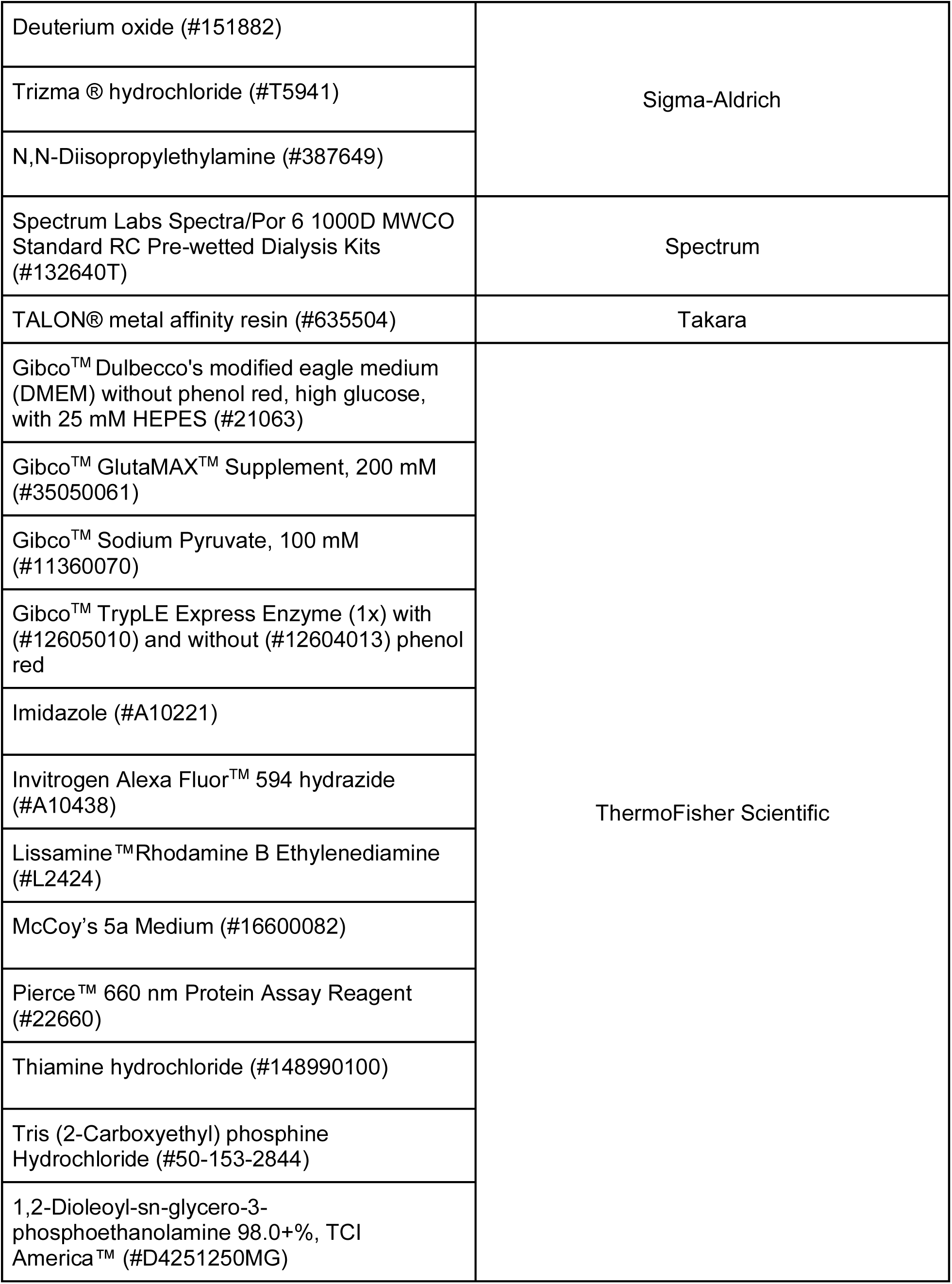
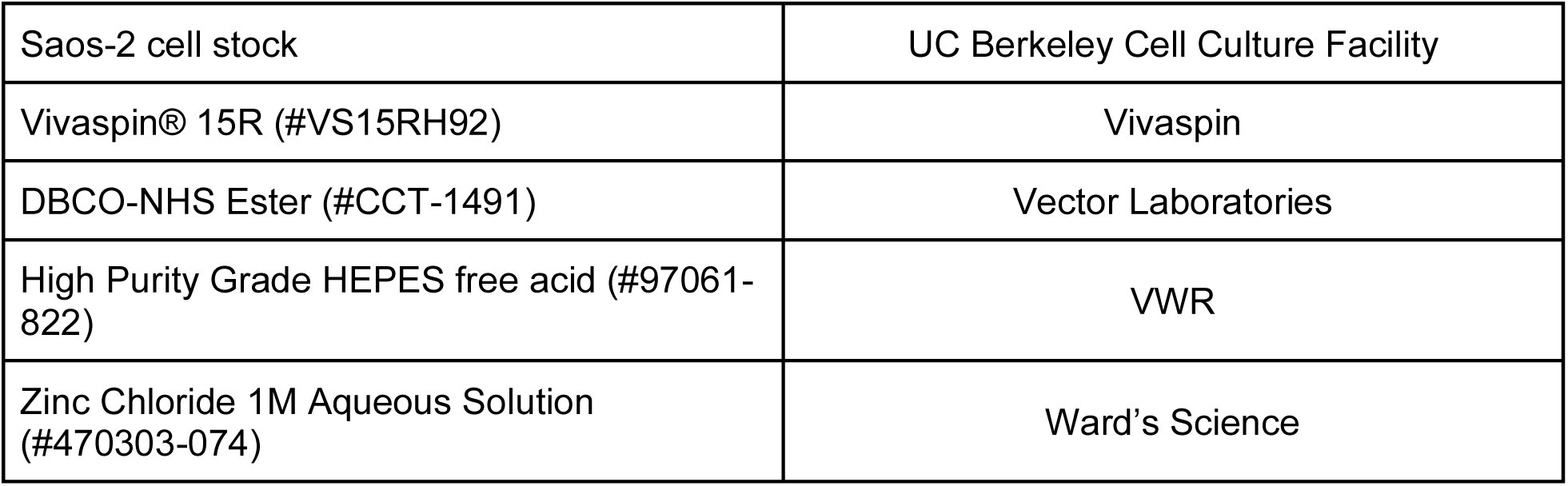
Materials.

**Table Supplemental Data Table 2 (S2).**
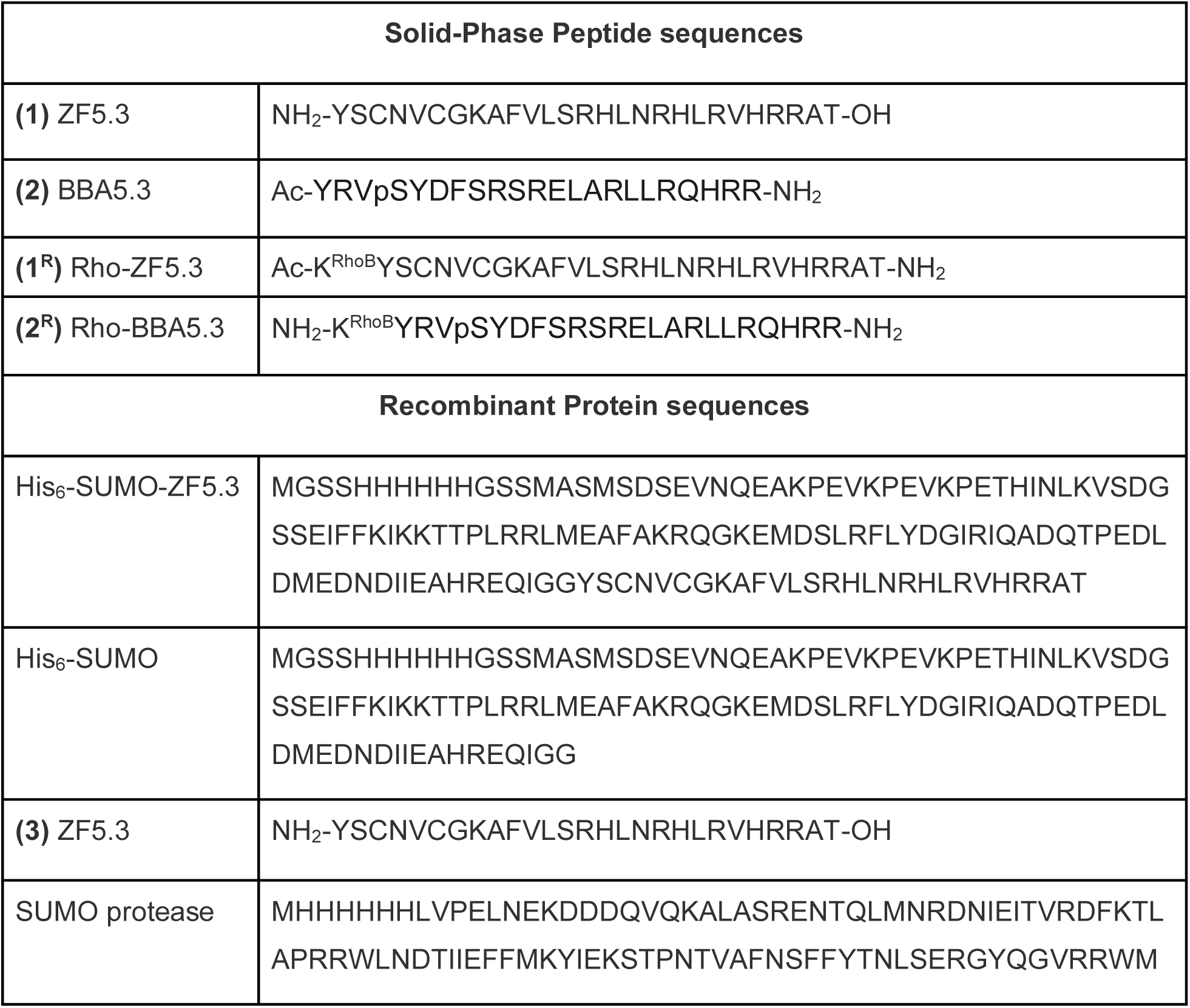
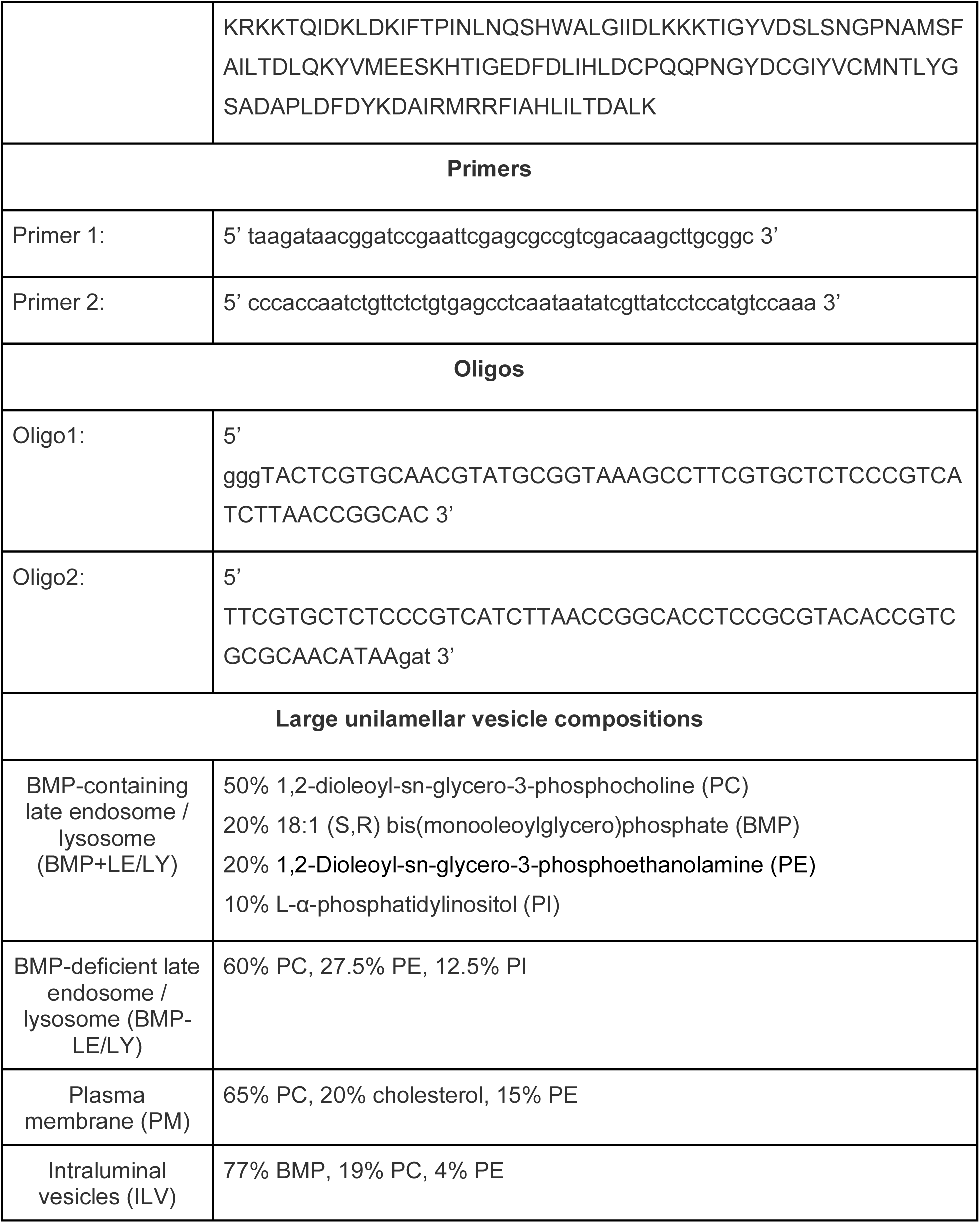
Relevant peptide, protein, DNA/RNA sequences, and large unilamellar vesicle compositions.

**Table Supplemental Data Table 3 (S3).** Relevant buffers.

| Buffer name | Composition |
| --- | --- |
| Lysis buffer: | 50 mM HEPES-HCl, 500 mM NaCl, 5 mM Imidazole, pH 7.4 |
| Wash Buffer: | 50 mM HEPES-HCl, 500 mM NaCl, 25 mM Imidazole, pH 7.4 |
| Elution Buffer: | 50 mM HEPES-HCl, 500 mM NaCl, 500 mM Imidazole, pH 7.4 |
| Dialysis Buffer: | 50 mM HEPES-HCl, 150 mM NaCl, 1 mM TCEP, pH 7.4 |
| Cation Exchange Buffer A: | 50 mM HEPES-HCl, 150 M NaCl, 1 mM TCEP, pH 7.4 |
| Cation Exchange Buffer B: | 50 mM HEPES-HCl, 2 M NaCl, 1 mM TCEP, pH 7.4 |
| Size Exclusion Buffer: | 50 mM Na<sub>2</sub>HPO<sub>4</sub>, 20 mM KH<sub>2</sub>PO<sub>4</sub>, and 9 mM NaCl pH 7.4 |
| Resuspension Buffer: | 20 mM Tris-HCl, 150 mM KCl, 1 mM TCEP, 100 uM ZnCl<sub>2</sub> pH 7.5 |
| Resuspension Buffer no Zn (II): | 20 mM Tris-HCl, 150 mM KCl, 1 mM TCEP, pH 7.5 |
| <sup>15</sup>N-<sup>1</sup>H HSQC Buffer: | 20 mM Citrate-phosphate, 100 mM NaCl, 2 mM TCEP, and 1.6 mM ZnCl<sub>2</sub> + 10% D<sub>2</sub>O |
| Structure determination Buffer: | 20 mM Tris-HCl, 100 mM NaCl, 2 mM TCEP, and 1.6 mM ZnCl<sub>2</sub> + 10% D<sub>2</sub>O |

**Table Supplemental Data Table 4 (S4).** Sample sizes for all intracellular delivery experiments.

| **Condition** | **Fig** | ***n* -FC** | **FC Mean** | **FC SEM** | ***n* -FCS** | **FCS Mean** | **FC SEM** |
| --- | --- | --- | --- | --- | --- | --- | --- |
| 1<sup>R</sup> - 1 μM, 1h | 5 | 2 | 4582 | 1311 | 29 | 146.4 | 21.68 |
| 2<sup>R</sup> - 1 μM, 1h | 5 | 3 | 1670 | 264.3 | 69 | 18.46 | 1.726 |
| 2<sup>R</sup> - 2 μM, 1h | 5 | 3 | 4796 | 1085 | 51 | 49.72 | 5.078 |
| 2<sup>R</sup> - 3 μM, 1h | 5 | 2 | 7545 | 767.0 | 16 | 46.20 | 5.007 |

**Table Supplemental Data Table 5 (S5).** Statistical information for all comparisons. All statistical tests were performed using GraphPad Prism 9 software. FCS data were analyzed using Brown-Forsythe and Welch ANOVA statistical tests followed by unpaired t-tests with Welch’s correction. The relevant parameters are defined below.

| Comparison | Mean Difference | 95.00% CI | P-value | Test statistic (t) | Degrees of freedom |
| --- | --- | --- | --- | --- | --- |
| **Figure 5f** |  |  |  |  |  |
| 1<sup>R</sup> - 1 μM, 1h vs<br>2<sup>R</sup> - 1 μM, 1h | 127.9 | 66.65 to 189.2 | <0.0001 | 5.882 | 28.36 |
| 1<sup>R</sup> - 1 μM, 1h vs<br>2<sup>R</sup> - 2 μM, 1h | 96.68 | 34.35 to 159.0 | 0.0008 | 4.341 | 31.10 |
| 1<sup>R</sup> - 1 μM, 1h vs<br>2<sup>R</sup> - 3 μM, 1h | 100.2 | 37.91 to 162.5 | 0.0005 | 4.503 | 30.90 |
| 2<sup>R</sup> - 1 μM, 1h vs<br>3<sup>R</sup> - 1 μM, 1h | 3.522 | -16.02 to 23.06 | 0.9968 | 0.4938 | 46.86 |
| **Figure 6a** |  |  |  |  |  |
| Comparison | Mean Difference | 95.00% CI | P-value | Test statistic (t) | Degrees of freedom |
| ZF5.3-ILV-pH4.5<br>vs pH7.5 | 0.1357 | 0.01297 to 0.2585 | 0.0229 | 3.227 | 32 |
| ZF5.3-BMP-LE/LY-pH4.5 vs pH7.5 | 0.02155 | -0.1012 to 0.1443 | 0.9995 | 0.5125 | 32 |
| ZF5.3-PM-pH4.5<br>vs pH7.5 | 0.07078 | -0.05196 to 0.1935 | 0.5776 | 1.683 | 32 |
| BBA5.3-ILV-pH4.5 vs pH7.5 | 0.1317 | 0.008958 to 0.2544 | 0.0293 | 3.131 | 32 |
| BBA5.3-BMP-LE/LY-pH4.5 vs<br>pH7.5 | 0.1177 | -0.005061 to 0.2404 | 0.067 | 2.798 | 32 |
| BBA5.3-PM-pH4.5<br>vs pH7.5 | 0.05944 | -0.06330 to 0.1822 | 0.7686 | 1.413 | 32 |
| BBA5.3-BMP+LE/LY-pH4.5<br>vs pH7.5 | 0.1355 | 0.01274 to 0.2582 | 0.0232 | 3.221 | 32 |
| ZF5.3-BMP+LE/LY-pH4.5 | 0.1339 | 0.01113 to 0.2566 | 0.0256 | 3.183 | 32 |

| vs pH7.5 |  |  |  |  |  |
| --- | --- | --- | --- | --- | --- |
| **Figure 6c** |  |  |  |  |  |
| **Comparison** | **Mean Difference** | **95.00% CI** | **P-value** | **Test statistic (t)** | **Degrees of freedom** |
| ZF5.3-ILV-pH7.5 vs pH4.5 | -0.0577 | -0.09220 to -0.02321 | 0.0003 | 7.093 | 15.51 |
| ZF5.3-BMP-LE/LY-pH7.5 vs pH4.5 | -0.3775 | -0.5833 to -0.1717 | 0.0002 | 8.091 | 12.93 |
| ZF5.3-PM-pH7.5 vs pH4.5 | 0.005066 | N/A | >0.9999 | 0.1766 | 8.216 |
| BBA5.3-ILV-pH7.5 vs pH4.5 | 0.004502 | N/A | >0.9999 | 0.4718 | 17.98 |
| BBA5.3-BMP-LE/LY-pH7.5 vs pH4.5 | -0.00235 | N/A | >0.9999 | 0.2404 | 15.13 |
| BBA5.3-PM-pH7.5 vs pH4.5 | -0.1235 | -0.2380 to -0.008985 | 0.0263 | 4.527 | 16.8 |
| ZF5.3-LE-pH7.5 vs pH4.5 | -0.04323 | -0.07766 to -0.008797 | 0.0064 | 5.323 | 15.52 |

## References cited

1. Bhardwaj, G. et al. Accurate de novo design of membrane-traversing macrocycles. Cell 185, 3520–3532.e26 (2022).

2. Doherty, G. J. & McMahon, H. T. Mechanisms of endocytosis. Annu. Rev. Biochem. 78, 857–902 (2009).

3. Mettlen, M., Chen, P.-H., Srinivasan, S., Danuser, G. & Schmid, S. L. Regulation of Clathrin-Mediated Endocytosis. Annu. Rev. Biochem. 87, 871–896 (2018).

4. Frankel, A. D. & Pabo, C. O. Cellular uptake of the tat protein from human immunodeficiency virus. Cell 55, 1189–1193 (1988).

5. Green, M. & Loewenstein, P. M. Autonomous functional domains of chemically synthesized human immunodeficiency virus tat trans-activator protein. Cell 55, 1179–1188 (1988).

6. Appelbaum, J. S. et al. Arginine topology controls escape of minimally cationic proteins from early endosomes to the cytoplasm. Chem. Biol. 19, 819–830 (2012).

7. LaRochelle, J. R., Cobb, G. B., Steinauer, A., Rhoades, E. & Schepartz, A. Fluorescence Correlation Spectroscopy Reveals Highly Efficient Cytosolic Delivery of Certain Penta-Arg Proteins and Stapled Peptides. J. Am. Chem. Soc. 137, 2536–2541 (2015).

8. Wissner, R. F., Steinauer, A., Knox, S. L., Thompson, A. D. & Schepartz, A. Fluorescence Correlation Spectroscopy Reveals Efficient Cytosolic Delivery of Protein Cargo by Cell-Permeant Miniature Proteins. ACS Cent. Sci. 4, 1379–1393 (2018).

9. Knox, S. L., Wissner, R., Piszkiewicz, S. & Schepartz, A. Cytosolic Delivery of Argininosuccinate Synthetase Using a Cell-Permeant Miniature Protein. ACS Cent. Sci. 7, 641–649 (2021).

10. Zoltek, M. et al. HOPS-Dependent Endosomal Escape Demands Protein Unfolding. ACS Cent. Sci. (2024) doi:10.1021/acscentsci.4c00016.

11. Zhang, X. et al. Dose-Dependent Nuclear Delivery and Transcriptional Repression with a Cell-Penetrant MeCP2. ACS Cent. Sci. 9, 277–288 (2023).

12. Shen, F. et al. A Cell-Permeant Nanobody-Based Degrader That Induces Fetal Hemoglobin. ACS Cent. Sci. 8, 1695–1703 (2022).

13. Steinauer, A. et al. HOPS-dependent endosomal fusion required for efficient cytosolic delivery of therapeutic peptides and small proteins. Proc. Natl. Acad. Sci. U. S. A. 116, 512– 521 (2019).

14. Mellman, I., Fuchs, R. & Helenius, A. Acidification of the endocytic and exocytic pathways. Annu. Rev. Biochem. 55, 663–700 (1986).

15. Krizek, B. A., Amann, B. T., Kilfoil, V. J., Merkle, D. L. & Berg, J. M. A consensus zinc finger peptide: design, high-affinity metal binding, a pH-dependent structure, and a His to Cys sequence variant. J. Am. Chem. Soc. 113, 4518–4523 (1991).

16. Bodenhausen, G. & Ruben, D. J. Natural abundance nitrogen-15 NMR by enhanced heteronuclear spectroscopy. Chem. Phys. Lett. 69, 185–189 (1980).

17. Urbani, A. et al. The Metal Binding Site of the Hepatitis C Virus NS3 Protease: A SPECTROSCOPIC INVESTIGATION*. J. Biol. Chem. 273, 18760–18769 (1998).

18. Pelton, J. G. et al. Structures of Active Site Histidine Mutants of IIIGlc, a Major Signal-transducing Protein in Escherichia coli: EFFECTS ON THE MECHANISMS OF REGULATION AND PHOSPHORYL TRANSFER*. J. Biol. Chem. 271, 33446–33456 (1996).

19. Cavanagh, J., Fairbrother, W. J., Palmer, A. G., Rance, M. & Skelton, N. J. CHAPTER 10-SEQUENTIAL ASSIGNMENT, STRUCTURE DETERMINATION, AND OTHER APPLICATIONS. in Protein NMR Spectroscopy (Second Edition) (eds. Cavanagh, J., Fairbrother, W. J., Palmer, A. G., Rance, M. & Skelton, N. J.) 781–817 (Academic Press, Burlington, 2007). doi:10.1016/B978-012164491-8/50012-9.

20. Grzesiek, S. & Bax, A. Correlating backbone amide and side chain resonances in larger proteins by multiple relayed triple resonance NMR. J. Am. Chem. Soc. 114, 6291–6293 (1992).

21. Skinner, S. P. et al. CcpNmr AnalysisAssign: a flexible platform for integrated NMR analysis. J. Biomol. NMR 66, 111–124 (2016).

22. Klukowski, P., Riek, R. & Güntert, P. Rapid protein assignments and structures from raw NMR spectra with the deep learning technique ARTINA. Nat. Commun. 13, 6151 (2022).

23. Klukowski, P., Riek, R. & Güntert, P. NMRtist: an online platform for automated biomolecular NMR spectra analysis. Bioinformatics 39, btad066 (2023).

24. Shen, Y. & Bax, A. Protein Structural Information Derived from NMR Chemical Shift with the Neural Network Program TALOS-N. in Artificial Neural Networks (ed. Cartwright, H.) 17–32 (Springer, New York, NY, 2015). doi:10.1007/978-1-4939-2239-0_2.

25. Schmidt, E. & Güntert, P. A New Algorithm for Reliable and General NMR Resonance Assignment. J. Am. Chem. Soc. 134, 12817–12829 (2012).

26. Güntert, P. Automated NMR structure calculation with CYANA. Methods Mol. Biol. Clifton NJ 278, 353–378 (2004).

27. Güntert, P. & Buchner, L. Combined automated NOE assignment and structure calculation with CYANA. J. Biomol. NMR 62, 453–471 (2015).

28. Schwieters, C. D., Kuszewski, J. J., Tjandra, N. & Clore, G. M. The Xplor-NIH NMR molecular structure determination package. J. Magn. Reson. San Diego Calif 1997 160, 65– 73 (2003).

29. Schwieters, C. D., Bermejo, G. A. & Clore, G. M. Xplor-NIH for molecular structure determination from NMR and other data sources. Protein Sci. Publ. Protein Soc. 27, 26–40 (2018).

30. Meng, E. C. et al. UCSF ChimeraX: Tools for structure building and analysis. Protein Sci. 32, e4792 (2023).

31. Williamson, M. P. Chemical Shift Perturbation. in Modern Magnetic Resonance (ed. Webb, G. A.) 995–1012 (Springer International Publishing, Cham, 2018). doi:10.1007/978-3-319-28388-3_76.

32. Vuister, G. W. & Bax, A. Quantitative J correlation: a new approach for measuring homonuclear three-bond J(HNH.alpha.) coupling constants in 15N-enriched proteins. J. Am. Chem. Soc. 115, 7772–7777 (1993).

33. Struthers, M., Ottesen, J. J. & Imperiali, B. Design and NMR analyses of compact, independently folded BBA motifs. Fold. Des. 3, 95–103 (1998).

34. Ingólfsson, H. I. et al. Lipid Organization of the Plasma Membrane. J. Am. Chem. Soc. 136, 14554–14559 (2014).

35. Kobayashi, T. et al. Separation and Characterization of Late Endosomal Membrane Domains *. J. Biol. Chem. 277, 32157–32164 (2002).

36. Chevallier, J. et al. Lysobisphosphatidic acid controls endosomal cholesterol levels. J. Biol. Chem. 283, 27871–27880 (2008).

37. Matsuo, H. et al. Role of LBPA and Alix in Multivesicular Liposome Formation and Endosome Organization. Science 303, 531–534 (2004).

38. Pattanakitsakul, S. et al. Association of Alix with Late Endosomal Lysobisphosphatidic Acid Is Important for Dengue Virus Infection in Human Endothelial Cells. J. Proteome Res. 9, 4640–4648 (2010).

39. Roth, S. L. & Whittaker, G. R. Promotion of vesicular stomatitis virus fusion by the endosome-specific phospholipid bis(monoacylglycero)phosphate (BMP). FEBS Lett. 585, 865–869 (2011).

40. Yang, S.-T., Zaitseva, E., Chernomordik, L. V. & Melikov, K. Cell-Penetrating Peptide Induces Leaky Fusion of Liposomes Containing Late Endosome-Specific Anionic Lipid. Biophys. J. 99, 2525–2533 (2010).

41. Brock, D. J. et al. Efficient cell delivery mediated by lipid-specific endosomal escape of supercharged branched peptides. Traffic 19, 421–435 (2018).

42. Larsson, E., Hubert, M. & Lundmark, R. Analysis of Protein and Lipid Interactions Using Liposome Co-sedimentation Assays. in Caveolae: Methods and Protocols (ed. Blouin, C. M.) 119–127 (Springer US, New York, NY, 2020). doi:10.1007/978-1-0716-0732-9_11.

43. Melo, A. M., Prieto, M. & Coutinho, A. Quantifying Lipid-Protein Interaction by Fluorescence Correlation Spectroscopy (FCS). in Fluorescence Spectroscopy and Microscopy: Methods and Protocols (eds. Engelborghs, Y. & Visser, A. J. W. G.) 575–595 (Humana Press, Totowa, NJ, 2014). doi:10.1007/978-1-62703-649-8_26.

44. Rhoades, E., Ramlall, T. F., Webb, W. W. & Eliezer, D. Quantification of α-Synuclein Binding to Lipid Vesicles Using Fluorescence Correlation Spectroscopy. Biophys. J. 90, 4692–4700 (2006).

45. Urbančič, I. et al. Lipid Composition but Not Curvature Is the Determinant Factor for the Low Molecular Mobility Observed on the Membrane of Virus-Like Vesicles. Viruses 10, 415 (2018).

46. Cymer, F., von Heijne, G. & White, S. H. Mechanisms of integral membrane protein insertion and folding. J. Mol. Biol. 427, 999–1022 (2015).

47. Ahlbach, C. L. et al. Beyond cyclosporine A: conformation-dependent passive membrane permeabilities of cyclic peptide natural products. Future Med. Chem. 7, 2121–2130 (2015).

48. Kino, T. et al. FK-506, a novel immunosuppressant isolated from a Streptomyces. II. Immunosuppressive effect of FK-506 in vitro. J. Antibiot. (Tokyo) 40, 1256–1265 (1987).

49. Kling, A. et al. Antibiotics. Targeting DnaN for tuberculosis therapy using novel griselimycins. Science 348, 1106–1112 (2015).

50. Bockus, A. T. et al. Probing the Physicochemical Boundaries of Cell Permeability and Oral Bioavailability in Lipophilic Macrocycles Inspired by Natural Products. J. Med. Chem. 58, 4581–4589 (2015).

51. van der Beek, J., de Heus, C., Sanza, P., Liv, N. & Klumperman, J. Loss of the HOPS complex disrupts early-to-late endosome transition, impairs endosomal recycling and induces accumulation of amphisomes. Mol. Biol. Cell 35, ar40 (2024).

52. Markley, J. L. et al. Recommendations for the presentation of NMR structures of proteins and nucleic acids (IUPAC Recommendations 1998). Pure Appl. Chem. 70, 117–142 (1998).

53. Findeisen, M., Brand, T. & Berger, S. A 1H-NMR thermometer suitable for cryoprobes. Magn. Reson. Chem. MRC 45, 175–178 (2007).

